# Extracellular matrix and microglial regulation of cognitive aging

**DOI:** 10.1101/2024.01.04.574215

**Authors:** Daniel T. Gray, Abigail Gutierrez, Yasaman Jami-Alahmadi, Vijaya Pandey, Lin Pan, Ye Zhang, James A. Wohlschlegel, Ross A. McDevitt, Lindsay M. De Biase

## Abstract

Synapse dysfunction is tightly linked to cognitive changes during aging. Emerging evidence suggests that microglia and the extracellular matrix (ECM) can potently regulate synapse integrity and plasticity. Yet the brain ECM, and its relationship with microglia, synapses, and cognition during aging remains virtually unexplored. Using ECM-optimized proteomic workflows and histological analyses, we discovered striking regional differences in ECM composition and aging-induced ECM remodeling across key basal ganglia nuclei. Moreover, we combine two distinct behavioral classification strategies with fixed-tissue confocal imaging and proteomic analysis to identify robust relationships between the hyaluronan- and proteoglycan-rich ECM and cognitive aging phenotypes. Finally, we provide evidence that aging midbrain microglia lose capacity to interact with and regulate the ECM, and that these aging-associated microglial changes are accompanied by local ECM accumulation and worse behavioral performance. Together, these foundational observations implicate changing microglia-ECM-synapse interactions as a key determinant of cognitive functioning during healthy aging.

## Main

Synapse loss and changes in synaptic plasticity are hallmark features of both normative brain aging and preclinical phases of neurodegenerative disease^1–3^. These synaptic changes have critical consequences, as preserved synapse status is linked to better cognitive outcomes in healthy aged rodents and nonhuman primates^1,4^. Growing evidence suggests the extracellular matrix (ECM) is a robust regulator of synaptic physiology during development and in early adulthood^5,6^. The ECM is not a static structure, but rather a dynamic network of proteins and carbohydrates remodeled to support numerous brain functions including synaptic plasticity and tissue repair^5–7^. Histochemical staining of ECM components suggests that ECM abundance^8^ and composition^9,10^ vary across brain regions and that ECM remodeling can occur during aging^7,11,12^. This raises the possibility that ECM status helps determine vulnerability of specific brain regions to aging-related synaptic decline.

Relative to other bodily tissues, the brain ECM is relatively depleted in fibrous ECM proteins like collagens and elastins and enriched in glycosylated proteins like proteoglycans^5,13,14^. Ongoing regulation of the brain ECM is a cooperative effort amongst different neuronal, vascular, and glial cell subtypes involved in ECM protein and carbohydrate synthesis, assembly, and degradation^5,6,15^. Microglia, the brain’s innate immune cells, express numerous ECM-relevant degradative enzymes as well as protease inhibitors^16–18^, positioning them as potent modifiers of ECM structure. Microglia also regulate synapses^19–24^, including through a recently discovered mechanism involving targeted degradation of ECM proteoglycans^7,25,26^. Moreover, microglial properties are substantially altered during aging, and changes in microglial density in the cerebral cortex are associated with ECM alterations in mouse models of Alzheimer’s disease and aging macaques^27,28^. Together, these observations suggest that changes in microglia-ECM interactions may play central roles in shaping synapse dynamics during aging, and consequently patterns of age-associated cognitive decline. Yet, this microglia-ECM-synapse triad remains virtually unexamined in most brain regions in both young adulthood and aging.

Numerous cognitive deficits associated with normative aging have been linked with dysregulation in midbrain dopaminergic circuits^29,30^. Microglia near midbrain dopamine neurons exhibit robust aging-related phenotypes characterized by increases in proliferation and inflammatory factor production that are evident by middle age in mice^31^. Whether these premature microglia aging phenotypes coincide with changes in ECM composition, and whether microglial-ECM responses to aging impact cognitive phenotypes remains unexamined. This study combines high-resolution imaging, quantitative tissue proteomics, and sophisticated behavioral-characterization strategies to comprehensively map how microglial-ECM responses to aging in the basal ganglia align with individual differences in synapse status and cognition.

## Results

### ECM-optimized proteomics reveals strong associations between ECM and synapse abundance in the aging basal ganglia

ECM proteins are highly glycosylated and comparatively insoluble, making it difficult to leverage traditional proteomic approaches to comprehensively map the brain ECM (i.e., matrisome). We compared two tissue processing workflows that have been used for ECM enrichment and analysis in peripheral bodily tissues: solubility-based subcellular fractionation^32^ and chaotropic extraction and digestion^33^. While the two approaches identified similar numbers of structural ECM proteins (core matrisome) in brain tissue, subcellular fractionation on average was more efficient at enriching ECM proteoglycans, while chaotropic digestion better enriched more fibrous ECM proteins like collagens **(Ext. Data Fig. 1**). Given the relative enrichment of proteoglycans in brain tissue^13,14^, we used the solubility-based workflow to create the first comprehensive proteomic mapping of the aging ECM proteome in the midbrain and striatum, two key basal ganglia nuclei, from young-adult (3 months) and aged (20+ months) wild-type mice (**Figure 1a**).

**Figure 1.**
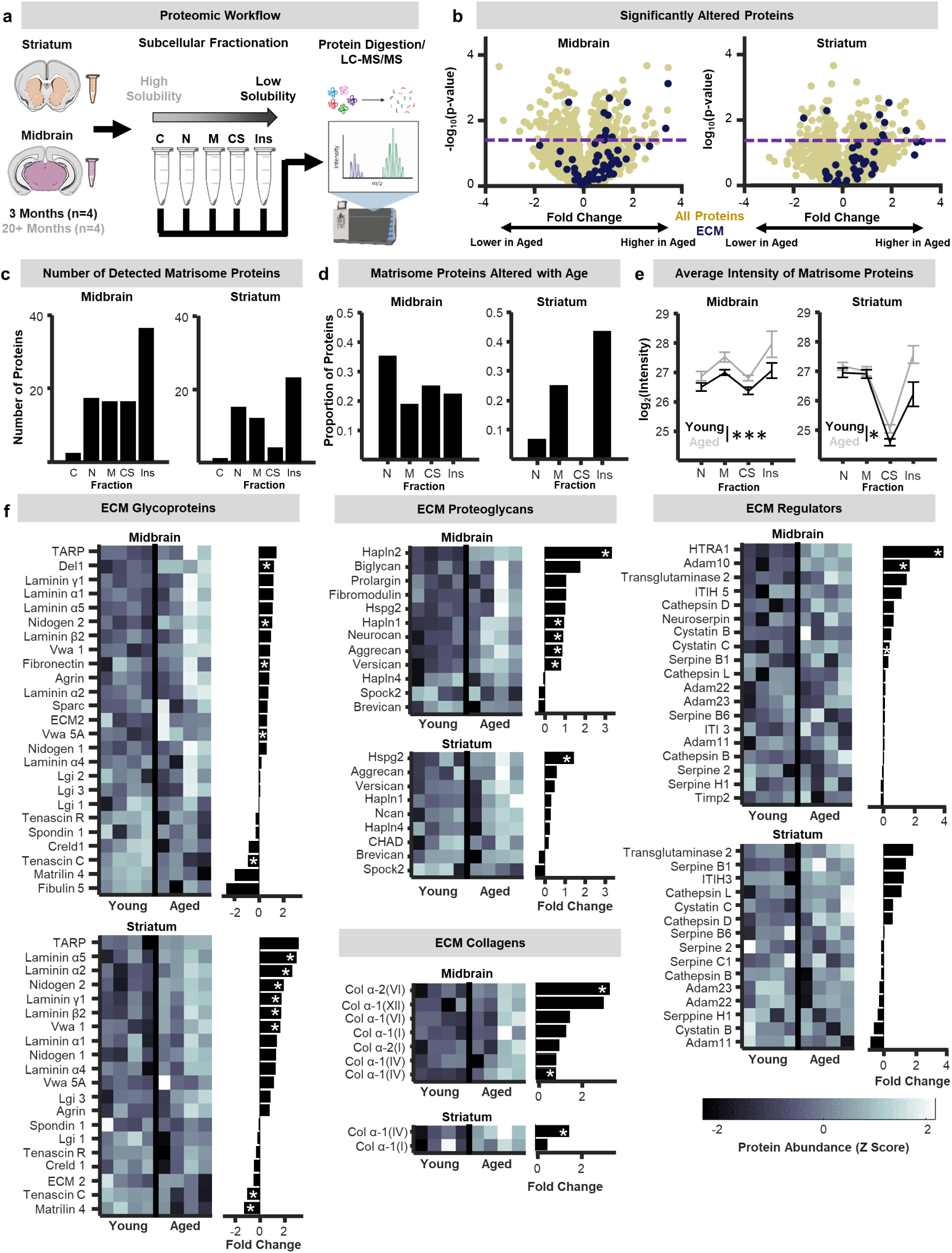
**a)** Schematic of the proteomic workflow. The midbrain and striatum of young-adult (3 months) and aged (20-24 months) wild-type mice were extracted. Tissue underwent a solubility-based subcellular fractionation protocol that yielded multiple samples of proteins with decreasing solubility that correspond to cytoplasmic, nuclear, membrane, and cytoskeletal subcellular localizations, and a final insoluble pellet. All samples were then sent for liquid chromatography with tandem mass spectrometry (LC-MS/MS) analysis. **b)** Volcano plots of all proteins pooled across solubility fractions for the midbrain and striatum. Dark blue dots denote individual structural extracellular matrix (ECM) proteins (e.g., core matrisome) detected in each region. Fold-changes were calculated with respect to the aged mice (positive values indicate greater with age). **c)** Bar plots of the number of core matrisome proteins detected across the different subcellular fractions in the midbrain and striatum. **d)** Bar plots depicting the proportion of structural ECM proteins that exhibited significant increases or decreases in abundance with age (p < 0.05; unpaired t-test). **e)** Average log2-transformed intensity values of core matrisome proteins across the nuclear, membrane, cytoskeletal, and insoluble fractions for young-adult (black) and aged (grey) mice in the midbrain and striatum. Protein concentrations were higher in the aged mice across the nuclear, membrane, cytoskeletal, and insoluble fractions in the midbrain (p < 0.001; n-way ANOVA) and striatum (p < p < 0.001; n-way ANOVA). **f)** Heat maps of protein abundances (z scored) and bar plots of corresponding fold-changes with age for ECM glycoproteins, proteoglycans, collagens, and ECM regulators (* p < 0.05; unpaired t-test).

Volcano plots of the abundance of all proteins in the tissue showed a relatively even split between proteins that were up- and down-regulated during aging (**Figure 1b**). In contrast, most core matrisome proteins increased in abundance during aging. Beyond protein abundance, changes to the structure and assembly of the matrix can occur and will be reflected in alterations to solubility of specific ECM components. As expected, in both brain regions, ECM proteins were primarily detected in more insoluble tissue fractions (**Figure 1c**). In the midbrain, individual ECM proteins exhibiting significant aging-related changes in abundance were relatively equally distributed across solubility fractions (**Figure 1d**). In the striatum, however, the majority of individual matrisome proteins that changed in abundance with age were found in the most insoluble fraction (**Figure 1d**). These regional differences were also observed at the level of protein abundances such that midbrain ECM protein abundances showed similar aging-related increases across subcellular fractions (**Figure 1e**) whereas striatum ECM protein abundances disproportionately increased within insoluble fractions during aging (**Figure 1e**). To probe this data further, we determined the number of ECM proteins showing significant changes in solubility during aging. This analysis revealed that just 6 percent of midbrain ECM proteins became more soluble in older animals and none more insoluble, whereas no striatal ECM protein became more soluble and 18 percent became more insoluble (**Ext. Data Fig 1**). These results indicate there are important regional differences in both the abundance and solubility of ECM proteins at different points of the lifespan.

Core matrisome proteins fall into 3 subclasses: glycoproteins, proteoglycans, and collagens. Comparable numbers of glycoproteins were detected in the midbrain and striatum, and several were significantly upregulated in both regions with aging (e.g., VWA 1, TARP). Laminin glycoproteins were specifically upregulated in the aging striatum (**Figure 1f**). More proteoglycans were detected in the midbrain compared to striatum, and nearly half of midbrain proteoglycans showed significant increases in abundance with age compared to just 1 in the striatum (**Figure 1f**). Collagen abundances also varied prominently between regions, with eight distinct collagens detected in the midbrain compared to just two in striatum (**Figure 1f**). Finally, several ECM regulatory proteins, which include metalloproteinases and protease inhibitors, were significantly more abundant in the aged midbrain (HTRA1, ADAM10, Cystatin C), whereas none changed in abundance with aging in the striatum (**Figure 1f**). Together, these observations demonstrate that regional heterogeneity in ECM composition and modulation during aging extends to all three subclasses of ECM proteins and their regulators.

To relate matrisome status with other features of the whole tissue proteome, Weighted Gene Coexpression Network Analysis^34^ was leveraged for unbiased identification of protein co-expression patterns (**Figure 2a**). This analysis identified 12 modules of covarying proteins. Most core matrisome (∼82%) and synapse proteins (∼63%) were members of yellow, brown, or tan modules (**Figure 2a**), highlighting these modules as warranting further analysis. Innate immune proteins, which can play key roles in synapse remodeling^23^, were also found within yellow (15%) and brown modules (12%), in addition to prominent presence in the turquoise (33%) module. Examination of module eigengenes revealed stark regional differences for the brown, yellow, and tan modules, but not the turquoise module (**Figure 2b**). Pathway analysis indicated that brown module proteins were associated with numerous metabolic processes, yellow with chemical synaptic transmission, turquoise with RNA metabolism and translation, and tan with a mix of biological processes including ECM organization (**Figure 2c**).

**Figure 2.**
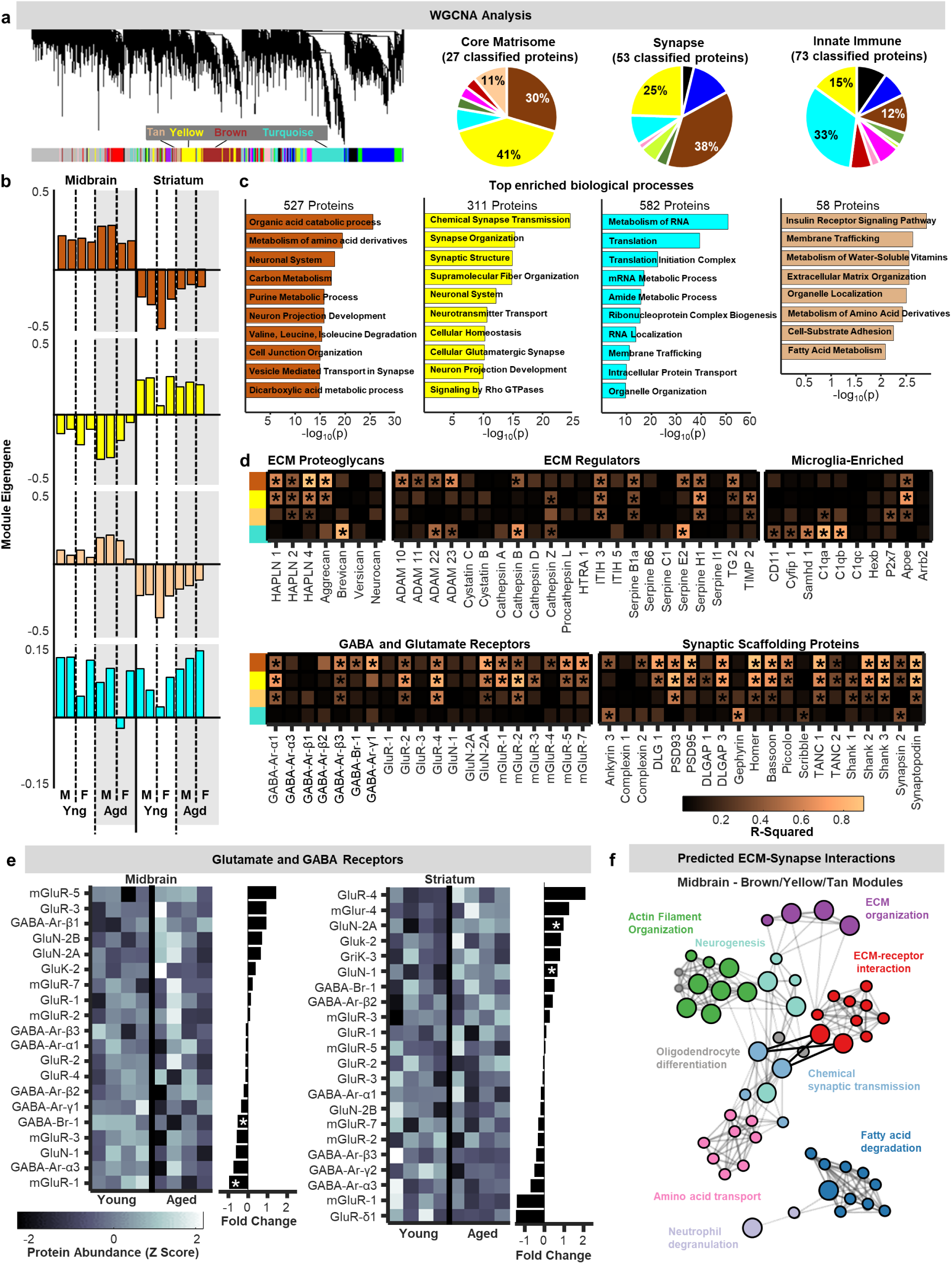
**a)** Left: dendrogram generated by Weighted Gene Coexpression Network Analysis (WGCNA) of all midbrain and striatal proteomic samples. Right: pie charts of classified core matrisome, synapse, and innate immune protein distributions across modules. **b)** Module eigengenes for the brown, yellow, tan, and turquoise modules for each sample. One aged female was classified as an outlier by the WGCNA and was removed from subsequent analyses. **c)** Enriched biological terms associated with brown, yellow, tan, and turquoise module proteins. **d)** Heatmaps representing correlations (r^2^ values; linear regression) between brown, yellow, tan, and turquoise module eigengenes and abundances of specific extracellular matrix proteoglycans, complement proteins, GABAergic receptors, glutamatergic receptors, and synaptic scaffolding proteins. **e)** Heat maps of z-scored protein abundances and bar plots of corresponding fold-changes with age for all detected glutamate and GABA receptors (* p < 0.05; unpaired t-test). **f)** Network plots of process and pathway enrichment terms associated with all midbrain tan/yellow/brown module proteins whose abundances were modulated by age (p < 0.05; Fisher’s exact test).

To further evaluate relationships between these modules and ECM, synapse, and immune proteins, we treated individual protein abundances as “traits” and examined their correlation with module eigengenes across samples (**Figure 2d**). Striking associations were observed between ECM proteoglycans and brown/yellow/tan module eigengenes, with the hyaluronan linker proteins HAPLN1-4 and aggrecan showing significant correlations. Although many immune signaling proteins also showed significant associations with brown/yellow/tan module eigengenes (**Ext. Data Fig. 2**), complement proteins C1qA and C1qB, which are known to tag synapses for microglial engulfment^23^, were only significantly correlated with turquoise module eigengenes. Finally, numerous synaptic proteins showed relationships with brown/yellow/tan module eigengenes, whereas very few correlated significantly with the turquoise module. Collectively, these observations suggest that synaptic and ECM protein abundances are tightly linked in the basal ganglia, and that complement signaling may not be involved in targeted ECM remodeling in these brain regions.

Overall abundance of most synaptic proteins did not significantly differ between young-adult and aged mice, indicating that substantial synapse *loss* is likely not occurring in the basal ganglia during healthy aging (**Figure 2e**). This aligns with previous reports showing that, while synapse loss does occur in vulnerable brain regions during healthy aging, synapse numbers remain stable in many others^1,35^. To further explore relationships between age-associated changes in ECM composition (**Figure 1**) and synapse status, we carried out additional pathway analysis focusing only on module proteins that were significantly altered during aging. Visualizing identified pathways as network plots revealed interconnected clusters of brown/yellow/tan module proteins associated with ECM organization and ECM-receptor interactions in the midbrain. This cluster was directly connected to clusters associated with synapse organization (bolded lines; **Figure 2f**), meaning that functional annotation predicts that proteins within these pathway “nodes” influence one another. While similar ECM clusters were observed in the striatum, they were not directly connected to pathway nodes associated with synapse organization (**Ext. Data Fig. 2**), suggesting that age-associated ECM remodeling has region-specific relationships with synapse status. Together, this proteomic mapping of matrisome, synapse, and immune proteins indicates that, in a normative aging context, regional variation in ECM status plays key roles in establishing and/or maintaining regional basal ganglia synapse profiles.

### Mesolimbic ECM networks differ across region and age and are positioned for synapse interactions

Although proteomic approaches provide unbiased and comprehensive quantification of ECM protein abundance, they cannot reveal the morphology and spatial distribution of ECM components. For independent analysis of brain ECM proteoglycans during aging, we histologically examined the ventral tegmental area (VTA, midbrain) and nucleus accumbens (NAc, ventral striatum), first using Wisteria floribunda agglutinin (WFA), a lectin that preferentially labels N-acetylgalactosamine residues that are most commonly found on chondroitin sulfate proteoglycans (i.e., aggrecan, brevican, neurocan, phosphacan)^36–39^. High-resolution (63x) confocal images were acquired from young-adult (4 months) and late-middle-aged (18 months), wild-type C57Bl6 mice, and ECM field-of-view coverage was calculated. In both the VTA and NAc, WFA label was distributed relatively evenly across fields of view and perineuronal net accumulations were only occasionally observed, indicating presence of a prominent interstitial matrix in both regions (**Figure 3b**). WFA tissue coverage was similar between the VTA and NAc of young-adult mice, and substantially greater in the VTA of middle-aged mice compared to young (**Figure 3c**). Chondroitin sulfate proteoglycans are sometimes referred to as ‘hyalectans’ due to their ability to interact with the ubiquitous ECM scaffold hyaluronan via several different linker proteins (hyaluronan and proteoglycan link proteins; HAPLNs)^40^. Because proteomic mapping suggested that numerous hyalectans and HAPLNs were significantly upregulated in the aged midbrain (**Figure 1**), and because hyaluronan is positioned to play central roles in determining overall ECM tissue topology^26,41^, we also histologically examined hyaluronan in young-adult and late-middle-aged mice. Hyaluronan tissue coverage was greater in the VTA compared to NAc, and on average higher in the VTA during aging (**Figure 3e**). Together, these histological findings are consistent with proteomic detection of greater midbrain ECM protein levels during aging (**Figure 1f**) and indicate that one prominent feature of VTA aging is an accumulation of the proteoglycan-rich ECM.

**Figure 3.**
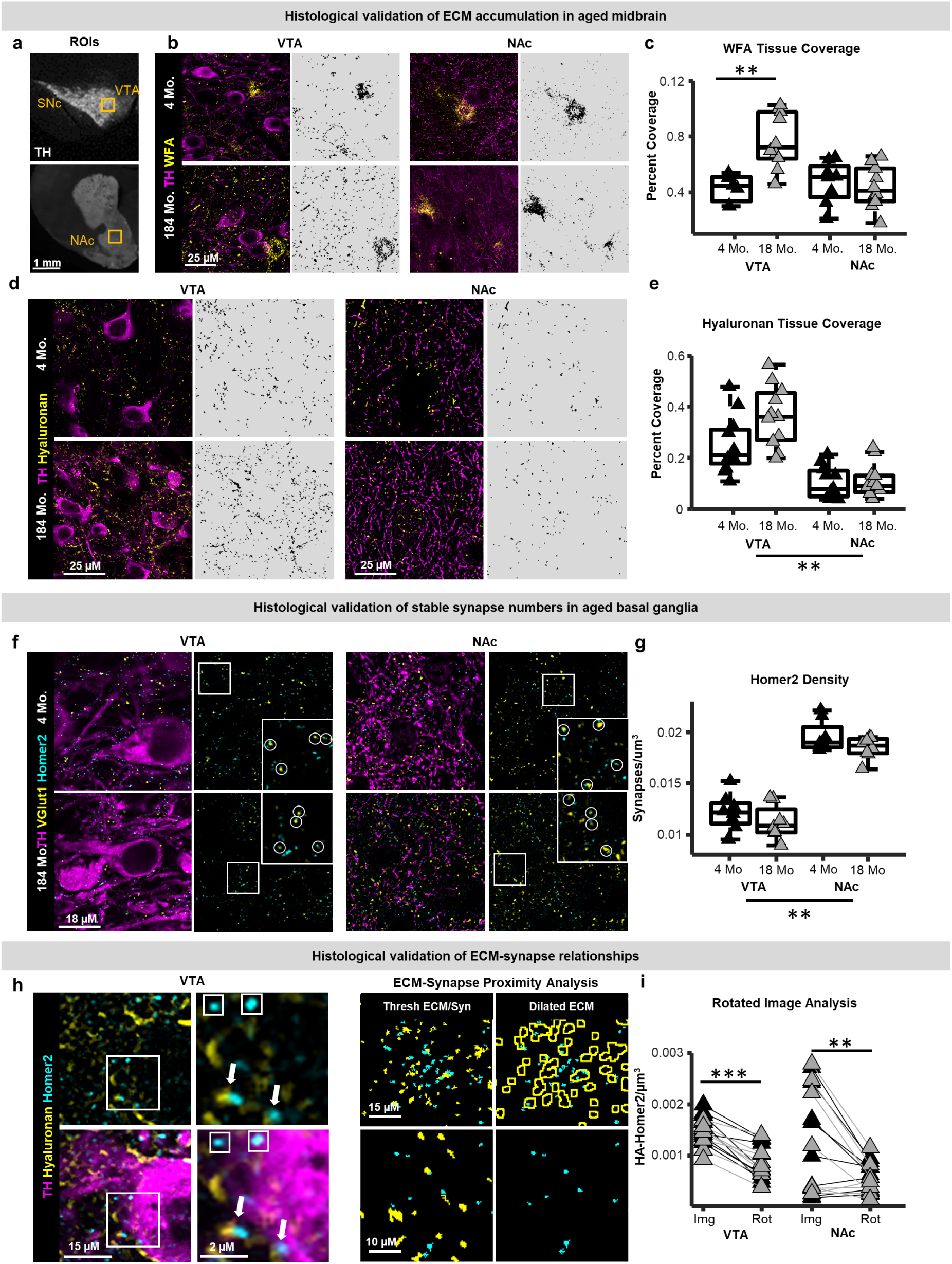
**a)** Histological validation of proteomic analysis of extracellular matrix (ECM) proteins was obtained from the ventral tegmental area (VTA; midbrain) and nucleus accumbens (NAc; striatum). **b)** Example photomicrographs of histochemically labelled tyrosine hydroxilase (TH) and Wisteria floribunda agglutinin (WFA) from a young-adult (4 months) and late-middle-aged (18 months) mouse (left panels). The right panels depict binarized images of the WFA matrix used for quantification. **c)** Boxplots depicting the proportion of a field of view occupied by WFA in the VTA and NAc of young-adult (black) and late-middle-aged (grey) mice. WFA tissue coverage was greater in the VTA of 18-month mice compared to young (p < 0.01; n-way ANOVA with post-hoc Tukey-Kramer) **d)** The ECM scaffold hyaluronan was analyzed in the same regions. The left panels show example photomicrographs of histochemically labelled hyaluronan and tyrosine hydroxylase in the VTA and NAc of a young-adult (4 months) and late-middle-aged (18 months) mouse (left panels). The right panels depict binarized images of the hyaluronan matrix used for quantification. **e)** Boxplots depicting the proportion of a field of view occupied by hyaluronan fibrils in the VTA and NAc of young-adult (black) and late-middle-aged (grey) mice. Hyaluronan tissue coverage was greater in the VTA compared to NAc (p < 0.01; n-way ANOVA), and on average greater in the VTA of late-middle-aged mice compared to young-adults (p > 0.05; post-hoc Tukey-Kramer). **f)** Example photomicrographs of immunohistochemically labelled Homer2 (excitatory postsynaptic puncta), VGlut1 (excitatory presynaptic puncta), and tyrosine hydroxylase in the VTA and NAc of young-adult and middle-aged wild-type mice. Inset images on the right depict the field of view outlined by the white squares within each image. Circles within the inset images delineate synapses with both a presynaptic and postsynaptic element. **g)** Homer2 densities were greater in the NAc compared to the VTA (p < 0.01; n-way ANOVA), but not different with age in either region. **h)** Left panels: example photomicrographs of histochemically labelled hyaluronan, homer2, and tyrosine hydroxylase in the VTA of a young-adult wild-type mouse. Images in the bottom panel are the same as in the top panel with the addition of the tyrosine hydroxylase channel. Zoomed in images are of the fields of view depicted by the white squares on the left images. Arrows highlight putative hyaluronan-homer2 colocalized puncta and small squares depict homer2 puncta not associated with hyaluronan. Right panel: schematic ECM-synapse proximity analysis to identify synaptic puncta within <0.5 microns of ECM. **i)** Densities of homer2 within 0.5 µm of hyaluronan were significantly lower in images where 1 channel was rotated by 90 degrees in both the VTA (p < 0.0001; paired t-test) and NAc (p < 0.01; paired t-test).

Proteoglycans anchored to hyaluronan can impact synapses via multiple mechanisms, including limiting structural remodeling and regulating lateral diffusion of neurotransmitter receptors^7,42^. To relate ECM structure to local synapse status, densities of excitatory pre- and post-synaptic proteins (VGlut1 and Homer2, respectively) were quantified (**Figure 3f**). Both VGlut1 and Homer2 densities were greater in the NAc compared to VTA but not altered by aging in either region (**Figure 3g; Ext. Data Fig. 3)**, consistent with proteomic data suggesting minimal synapse loss in the aging midbrain and striatum. The abundance of colocalized VGlut1-Homer2 puncta, which may better represent functional synapses, also did not differ with age in either region (**Ext. Data Fig. 3).** To probe spatial relationships between ECM and synapses, hyaluronan fibrils were reconstructed and dilated by 0.5 μm, to estimate the density of Homer2 puncta within the potential territory of proteoglycans anchored to this scaffold (**Figure 3h**). In both regions, hyaluronan-homer2 spatial associations were greater than would be expected by chance (associations detected when rotating one fluorescence channel by 90 degrees, **Figure 3i**), supporting the idea of functional associations between local ECM and synapses. The density and proportion of Homer2 within 0.5 μm of hyaluronan was similar in young-adult and late-middle-aged mice and not different across regions, although greater variability was observed in the NAc. Together, these histological findings validate regional ECM heterogeneity revealed by proteomics, confirm regional differences in age-associated ECM remodeling, and suggest that ECM-synapse spatial associations occur throughout life.

### Microglia-ECM relationships during normative aging

VTA microglia exhibit robust aging-related phenotypes characterized by changes in proliferation, morphology, and inflammatory factor production^31^, raising the possibility that microglia contribute to ECM accumulations in the aging VTA. Furthermore, aging-related increases in ECM protein abundance at the level of tissue proteomics (**Figure 1**) arose alongside increases in the abundance of most detected microglia-enriched proteins with advanced age (**Figure 4a**). To examine whether changes in microglial and ECM abundance align in the aging VTA, we histologically co-labelled the ECM (hyaluronan) and microglia (IBA1) in young-adult (4 months) and late-middle-aged (18 months) wild-type mice and examined relationships between the two. VTA microglia densities were greater in middle-aged mice compared to young-adults (**Figure 4b**; see scatter plot). Microglia densities were significantly positively correlated with hyaluronan tissue coverage in young-adult mice, and this significant relationship was lost in the older mice (**Figure 4b**). Next, microglial morphology was evaluated using a 2-dimensional Sholl analysis and, as shown previously^31^, aged VTA microglia exhibited fewer Sholl intersections compared to young, indicative of a less complex morphology (**Ext. Data Fig. 4**). As with microglia densities, microglial morphological complexity was significantly positively correlated with hyaluronan deposition in young-adult mice and this relationship was lost in the older animals (**Figure 4c**). To determine whether VTA microglia make direct contact with the hyaluronan matrix, reconstruction of microglia and hyaluronan was carried out in Imaris using tissue from young-adult (3-4 months), middle-aged (12-17 months) and aged (18-22 months) Cx3Cr1^EGFP/+^ mice, which enable precise visualization of microglial morphology. VTA microglia in young mice made relatively regular putative contacts with hyaluronan fragments, both along their processes and proximal to microglial somas. Quantification of the density of hyaluronan contacts normalized to GFP signal revealed an age-associated decrease in microglia-hyaluronan contacts in the VTA (**Figure 4d**). This raises the intriguing possibility that loss of ability of VTA microglia to interact directly with hyaluronan networks is related in some way to greater hyaluronan deposition within the tissue.

**Figure 4.**
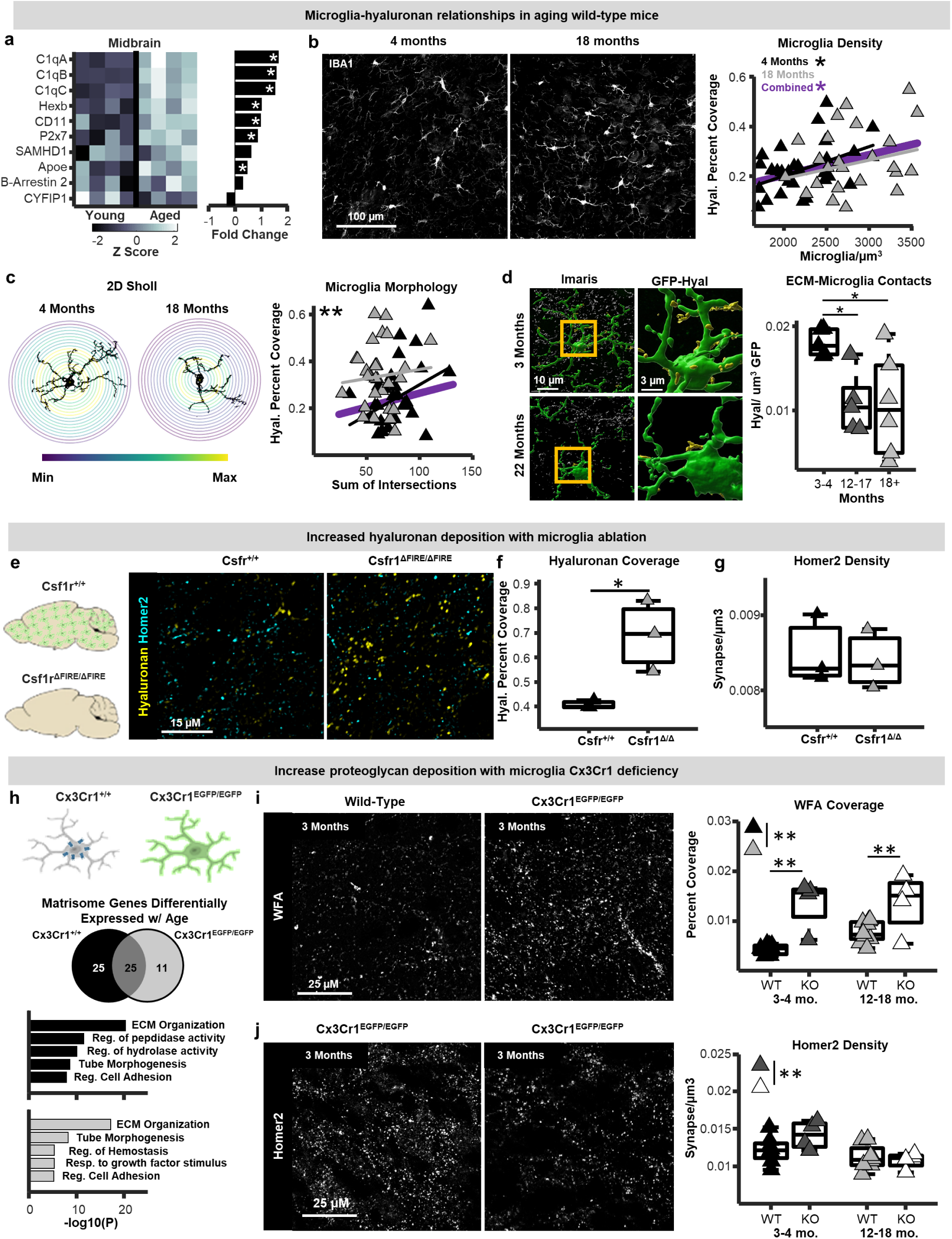
**a)** Heat maps of z-scored protein abundances and bar plots of corresponding fold-changes with age for all detected microglia-enriched proteins within the proteomic dataset from the young-adult and middle-aged midbrain (* p < 0.05; unpaired t-test). **b)** Left: example photomicrographs of IBA1-positive microglia in the ventral tegmental area (VTA) of young-adult (4 months) and late-middle-aged (18 months) wild-type mice. Right: scatter plots of the relationship between microglia densities and hyaluronan tissue coverage in the VTA. Microglia densities showed significant relationships with hyaluronan coverage in young-adult mice (p < 0.05; robust regression) that were absent in middle-aged mice. **c)** Microglia morphological complexity was assessed in wild-type mice using a 2-dimensional Sholl analysis of IBA1-positive VTA microglia. Left: example binarized VTA microglia and Sholl radii from 4-month and 18-month-old mice. Right: scatter plots of the relationship between microglia morphological complexities and hyaluronan tissue coverage in the VTA. Microglial complexities showed significant relationships with hyaluronan coverage in young-adult mice (p < 0.01; robust regression) that were absent in middle-aged mice. **e)** Hyaluronan matrix deposition and Homer2 densities were assessed in young-adult (3 months) Csf1r^ΔFIRE/ΔFIRE^ mice in which microglia are constitutively depleted, and Csf1r^+/+^ control mice. **f)** Hyaluronan tissue coverage and **g)** Homer2 puncta densities in the VTA of Csf1r^+/+^ (black triangles) and Csf1r^ΔFIRE/ΔFIRE^ (grey triangles) mice. Hyaluronan coverage was greater in Csf1r^ΔFIRE/ΔFIRE^ mice compared to Csf1r^+/+^ (p < 0.05; unpaired t-test), whereas Homer2 puncta densities were not different between genotypes. **h)** Top: ECM and synapse abundances were also examined in young-adult (3-4 months) and early-middle-aged (12-15 months) Cx3Cr1-knockout (Cx3Cr1^EGFP/EGFP^) mice. Middle: pie chart of the number of differentially expressed matrisome-related genes between 12-month and 2-month wild-type and Cx3Cr1-knockout mice (from Gyoneva et al., 2019). Bottom: enriched biological pathways associated with differentially expressed matrisome genes with advanced age in wild-type and Cx3Cr1-knockout mice. **i)** Left: example photomicrographs of histochemically labelled Wisteria floribunda agglutinin (WFA; ECM) in the VTA of wild-type and Cx3Cr1-knockout mice. Right: Boxplots of WFA tissue coverage in wild type (black triangles; light grey triangles) and Cx3Cr1-knockout mice (dark grey triangles; white triangles). Cx3Cr1-knockout mice exhibited greater WFA matrix tissue coverage compared to wild type (p < 0.001; n-way ANOVA) and only wild-type mice exhibited age-related increases in WFA tissue coverage (p < 0.05; post-hoc Tukey-Kramer). **j)** Left: example photomicrographs of immunohistochemically labelled Homer2 (postsynaptic) in the VTA of 3-month and 12-month-old Cx3Cr1-knockout mice. Right: boxplots of Homer2 puncta densities in wild-type and Cx3Cr1-knockout mice. Cx3Cr1-knockout mice exhibited a decrease in Homer2 densities with advanced age that the wild-type mice did not (p < 0.05; n-way ANOVA with post-hoc Tukey-Kramer).

### Young-adult VTA microglia attenuate ECM deposition

To probe observed correlations between ECM deposition and microglial properties in the VTA, we examined tissue from young-adult (3-4 months) microglia-deficient mice (Csf1r^ΔFIRE/ΔFIRE^ mice; **Figure 4e**)^43^. Compared to control mice, Csf1r^ΔFIRE/ΔFIRE^ mice exhibited higher VTA hyaluronan deposition but no difference in Homer2 densities (**Figure 4f, 4g**), as observed in aging wild-type mice (**Figure 3**). Together with findings that the ECM accumulates with age in the VTA (**Figure 3**), these observations suggest that young-adult microglia attenuate ECM deposition in the VTA and that this microglial function is diminished with normative aging. While elevated hyaluronan deposition was not associated with altered numbers of postsynaptic structures in either aging wild-type or microglia-deficient mice, these changes likely impact proteoglycan deposition around synapses and modify capacity for synapse plasticity^5^.

Next, we sought to examine the ECM in a context where microglia are present but undergoing distinct aging trajectories. Cx3Cr1 deficiency has been suggested to alter microglial aging phenotypes^31,44^, impact synaptic plasticity^45^, and regulate ECM composition in other bodily tissues^46,47^. Indeed, when we mined a published RNAseq dataset^44^ of microglia from young-adult (2 months) and middle-aged (12 months) wild-type (WT) and Cx3Cr1-deficient (KO) mice, we found that over 50 matrisome-related genes (both structural ECM proteins and ECM regulatory proteins) were differentially expressed in 2mo KO microglia compared to 2mo WT microglia (**Ext. Data Fig. 4**). Critically, more ECM-relevant genes were altered during aging in WT microglia compared to the number that were altered during aging in KO microglia, indicating that Cx3Cr1-deficiency perturbs ECM-related aspects of microglial aging. The most robust difference between genotypes was an age-related upregulation of genes associated with negative regulation of peptidase and hydrolase activity in WT microglia that was absent in KO microglia (**Figure 4h; Ext. Data Fig. 4**). Altogether, these results indicate that Cx3Cr1-deficiency is a suitable manipulation to probe how altered microglial aging trajectories impact the ECM.

Via immunostaining, we found that VTA microglia from KO (*Cx3Cr1^EGFP/EGFP^;* KO) mice exhibited reduced morphological complexity compared to Cx3Cr1-heterozygous (*Cx3Cr1^EGFP/+^,* HET) mice, confirming that this manipulation enhances features of VTA microglial aging that we have reported previously (**Ext. Data Fig. 4**)^31^. We then examined ECM (WFA) and synapse abundance (Homer2) in the VTA of young-adult (3-4 months) and middle-aged (12-18 months) WT and KO mice. WFA tissue coverage was significantly greater in KO mice compared to WT both in young-adulthood and middle-age (**Figure 4i**), and this same effect was also observed in the NAc (**Ext. Data Fig. 5**). Moreover, age-related ECM accumulations observed in WT mice was absent in KO mice (**Figure 3**; **Figure 4i**), further supporting the hypothesis that Cx3Cr1-defiency alters ECM-related aspects of microglial aging phenotypes. Intriguingly, hyaluronan abundance was not significantly different between WT and KO mice at any age (**Ext. Data. Fig. 5**), suggesting that effects of Cx3Cr1-deficiency on ECM regulation preferentially impact some ECM components over others. KO mice exhibited significant reductions in Homer2 density by middle age, unlike WT mice where synapse numbers were stable into late middle age (**Figure 3**; **Figure 4j**). Analysis of colocalization between microglia, hyaluronan, and homer2 in KO mice indicated that putative microglial hyaluronan engulfment was not different with age (**Ext. Data. Fig. 5**). Putative microglial synapse engulfment was also not different in young-adult and middle-aged KO mice (**Ext. Data. Fig. 5**), indicating that age-associated reductions in synapse density in these mice do not arise from greater microglial synapse engulfment. Together these results indicate that WT microglia have an attenuating effect on ECM deposition relative to contexts where microglia are absent or altered, and that during normative aging this microglial function is lost to some degree. It will be critical to replicate these observations using conditional microglial manipulations, as some findings may be influenced by developmental compensations that arise in constitutive models.

### Hyaluronan and synapse remodeling in the VTA aligns with reward-based memory in middle-aged mice

Healthy brain aging is an active process that engages endogenous mechanisms of plasticity to protect circuit function as aging-related challenges emerge^48^. To begin understanding whether ECM accumulations in the aging VTA represents vulnerabilities or adaptive responses that support continued circuit function, we developed a behavioral paradigm that engages reward circuitry and probes aspects of dopamine-relevant cognition known to be altered with aging, including reward memory and cognitive flexibility (**Figure 5a**)^29,49^. In this task, mice learn to explore a large arena and forage for palatable food reward that changes location daily. During testing, mice encode a rewarded location and, after 2 or 24 hours, reenter the arena with 4 unrewarded feeders to measure their memory of the rewarded location (probe trials; **Figure 5a**). Mice then immediately re-enter the arena and are allowed to consume food reward in the previously rewarded location to minimize extinction of training (**Figure 5a**). In total, mice undergo 5 weeks of training/testing (5 days/week), which allows for robust assessment of cognitive status of individual mice and minimizes behavioral variables like novelty and stress at the time when tissue is collected for histological examination.

**Figure 5.**
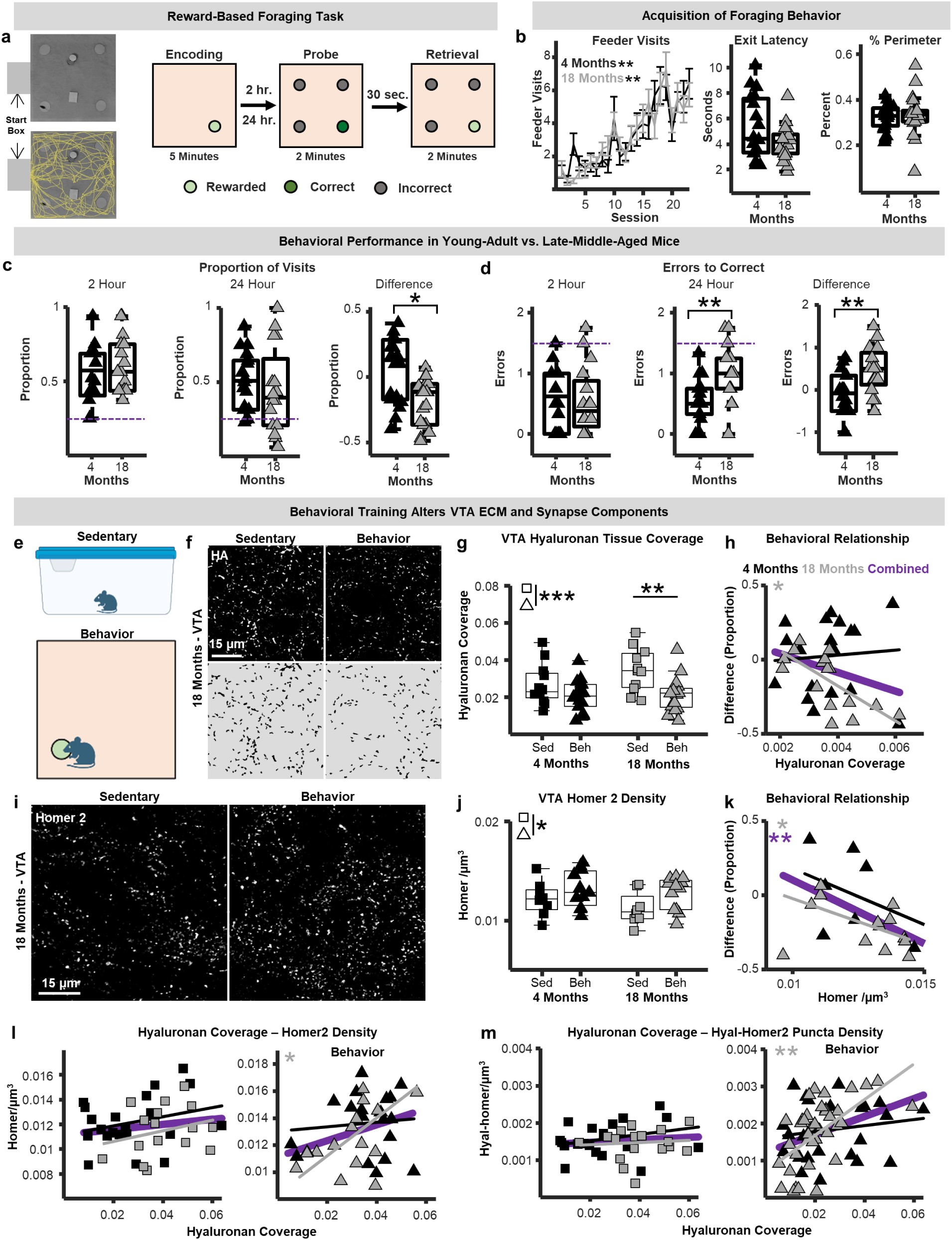
**a)** Left: schematic depicting the foraging arena with proximal (fixed objects within arena) reference landmarks, a start box equipped with a guillotine door, and overhead tracking of mouse behavior (yellow). Right: schematic depiction of the test phase of the reward-based foraging paradigm. **b)** Right: the number of feeder visits increased across sessions in both young-adult (4 months; black) and late-middle-aged (18 months; grey) mice (p < 0.01; repeated measures ANOVA). The latency to exit the start box (middle), and the proportion of time on the perimeter of the arena (right) were not different between age groups. **c)** The proportion of correct feeder visits during probe trials at 2- and 24-hour delays (left and middle), and the within-subject difference measure (24 hour-2 hour; right). Middle-aged mice made significantly fewer correct feeder visits at 24 hours compared to 2 hours (p < 0.05; n-way ANOVA), whereas the performance of the young-adult mice was similar between the two conditions. **d)** The number of errors to correct feeder visits during probe trials at 2- and 24-hour delays (left and middle), and the within subject difference measure (24 hr.-2 hr.). Middle-aged mice made more errors at the 24-hour condition compared to young (p < 0.01; unpaired t-test) and made significantly more errors at 24 hours compared to 2 hours. (p < 0.01; n-way ANOVA). **e)** Hyaluronan and Homer2 were assessed in the VTA of young-adult (4 months) and late-middle-aged (18 months) behavior-trained mice and sedentary controls. **f)** Top: example photomicrographs of histochemically labelled hyaluronan from a sedentary and behavior-trained 18-month-old mouse. Bottom: binarized images of the hyaluronan matrix used for quantification. **g)** Boxplots depicting VTA hyaluronan tissue coverage in young-adult (black) sedentary (squares) and behavior-trained (triangles) mice and late-middle-aged (grey) sedentary and behavior-trained mice. Behavior-trained mice exhibited lower hyaluronan coverage relative to controls (p < 0.001; n-way ANOVA), and behavior-trained middle-aged mice had lower hyaluronan levels relative to sedentary (p < 0.01; post-hoc Tukey-Kramer). **h)** Relationship between VTA hyaluronan coverage and performance on reward-based foraging task (proportion of correct feeder visits difference measure). A significant negative relationship was observed in middle-aged mice (p < 0.05; robust regression). **i)** Example photomicrographs of immunolabelled Homer2 from a sedentary and behavior-trained 18-month-old mouse. **j)** Boxplots of VTA Homer2 densities in young-adult sedentary and behavior-trained mice and late-middle-aged sedentary and behavior-trained mice. Behavior-trained mice exhibited higher Homer2 densities relative to controls (p < 0.05; n-way ANOVA). **k)** Relationship between VTA Homer2 densities and performance on reward-based foraging task. Significant negative relationships were observed across all mice (purple; p < 0.01; robust regression) and in middle-aged mice (p < 0.05; robust regression). **l)** Scatter plots of the relationship between hyaluronan tissue coverage and Homer2 puncta densities in young-adult and late-middle-aged sedentary and behavior-trained mice. A significant positive correlation was observed only in middle-aged behavior-trained mice (robust regression; r = 0.48; p < 0.05). **m)** Scatter plots of the relationship between hyaluronan tissue coverage and densities of Homer2 within 0.5 µm of hyaluronan fibrils in young-adult and late-middle-aged sedentary and behavior-trained mice. Again, a significant positive correlation was observed only in middle-aged behavior-trained mice (robust regression; r = 0.59; p < 0.01).

Both young-adult (4 months) and late-middle-aged (18 months) non-food-restricted mice learned the task and exhibited consistent foraging at similar points of the experiment timeline (**Figure 5b, Ext. Data Fig. 6**). Average walking speeds, start box exit latencies, and proportion of time on the arena perimeter did not differ between age groups, indicating similar levels of task engagement and absence of prominent age-related differences in anxiety-like behavior (**Figure 5b).** During probe sessions, young-adult mice showed similar performance that was better than chance following both 2- and 24-hour delays. Middle-aged mice performed comparably to young-adults with a 2-hour delay but made fewer correct feeder visits and more errors following 24-hour delays (**Figure 5c**). Critically, we observed higher variability in middle-aged mice during 24-hour probe trials compared to young, which is a hallmark of cognitive aging^50^. Hence, this nuanced behavioral paradigm establishes a strategy to probe links between cellular/molecular features of mesolimbic dopaminergic circuits and age-associated changes in reward-based cognition.

To enable assessment of the impact of the 5-week behavioral training itself, each cohort of mice included sedentary controls housed in the same vivarium as behaving mice for the duration of the experiment (**Figure 5e**). Compared to sedentary mice, behavior-trained mice exhibited significantly less hyaluronan within the VTA (**Figure 5f, 5g**), particularly at middle-age. Hence, the accumulation of VTA hyaluronan observed in aging sedentary animals appeared to be mitigated by engaging in behavioral training. Furthermore, middle-aged mice with lower hyaluronan densities showed better task performance (**Figure 5h**), indicating that this remodeling is beneficial. Behavior-induced hyaluronan remodeling was not observed in multiple other brain regions examined, including the NAc, mPFC, and retrosplenial cortex (**Ext. Data Fig. 6**). Behavior-associated hyaluronan reductions were observed in the substantia nigra pars compacta and hippocampus, suggesting that hyaluronan remodeling occurs only in specific circuits when animals repeatedly engage with reward-based spatial memory tasks.

To link behavior-induced VTA hyaluronan remodeling with synapse status, Homer2 was also analyzed in these mice. Compared to sedentary mice, behavior-trained mice had elevated Homer2 puncta densities, indicating that behavioral training/testing had net synaptogenic effects (**Figure 5i, 5j**). Surprisingly, however, behavior-trained mice with fewer synapses showed better task performance (**Figure 5k**), suggesting that synapse refinement that impacts performance may also occur across this 5-week paradigm. In sedentary mice, hyaluronan tissue coverage and Homer2 puncta densities were not correlated in either young-adult or late-middle-aged mice. Importantly, however, a significant positive correlation between hyaluronan tissue coverage and homer2 densities was observed in middle-aged mice that underwent behavioral training, but not in the young-adults (**Figure 5l**). Furthermore, behavior-trained middle-aged mice with more hyaluronan tissue coverage had a higher density of Homer2 puncta within 0.5 µm of hyaluronan (see ECM-synapse proximity analysis; **Figure 3h**), and again this relationship was not seen in young-adult mice or in sedentary mice at either age (**Figure 5m**). Together, these experiments suggest that greater VTA hyaluronan abundance may stabilize excitatory synapse numbers but limit synaptic refinements that optimize reward-driven behavior in aging mice.

### Proteomic signatures of cognitive phenotypes in middle-aged mice

Because histochemistry cannot provide comprehensive quantitative information on large families of proteins, we sought to use tissue proteomics to identify matrisome and synapse protein expression patterns associated with cognitive function in aging mice. To this end, young-adult (4 months) and late-middle-aged (18 months) wild-type mice were tested on 3 standard mouse behavioral paradigms: an open field test of anxiety-like behavior, a novel object recognition (NOR) test of non-spatial recognition memory, and a T-maze test of spontaneous alternation behavior (**Figure 6a**). These 3 behaviors were selected due to their relatively high-throughput nature, allowing for a more rapid assessment of cognitive status while minimizing remodeling effects of extended behavioral training, and because this behavioral battery will be more easily replicated across laboratories within different experimental contexts. Middle-aged mice on average exhibited lower discrimination and alternation on the NOR and T-maze tasks, respectively, both of which are indicative of poorer performance. As in the foraging paradigm (**Figure 5**), middle-aged mice also exhibited higher variability in performance across tasks. Leveraging this variability, we performed unbiased hierarchical clustering analysis to cognitively classify all mice tested in this pipeline. This analysis resulted in two parent clusters, one containing 75% of the young-adult mice and roughly half (46%) of the middle-aged mice, and another cluster containing the remaining animals. Using these clusters, we categorized the mice into 3 groups: young average, middle-aged unimpaired, and middle-aged impaired (**Figure 6a**). Midbrains were harvested from mice in each age and cognitive group and subjected to solubility-based subcellular fractionation (**Figure 1)**, and membrane, cytoskeletal, and insoluble fractions were analyzed via mass spectrometry.

**Figure 6.**
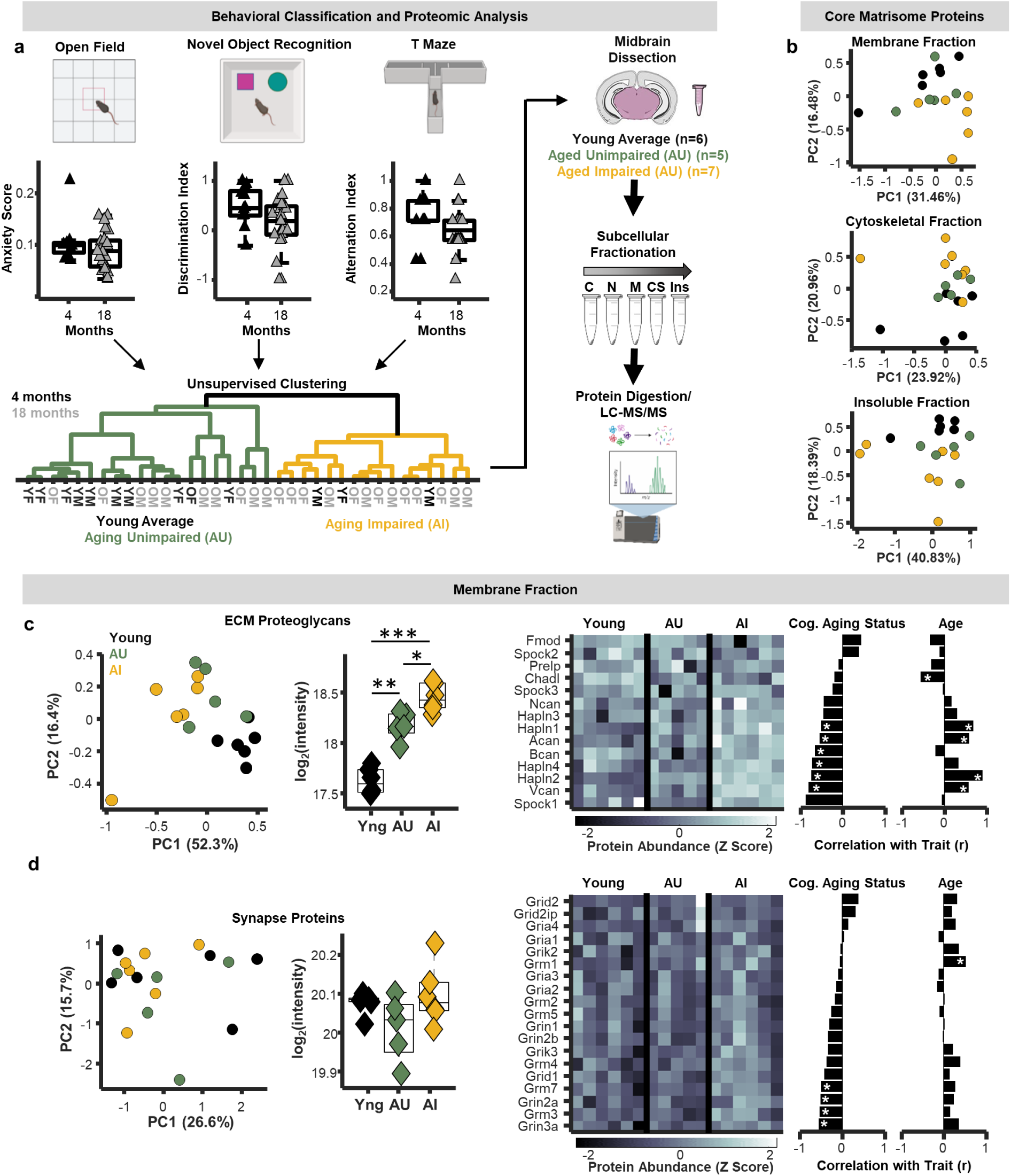
**a)** Schematic behavioral classification and proteomic analysis. Young-adult (4 months; n = 12; black triangles) and late-middle-aged (18 months; n = 24; grey triangles) male and female mice were tested on an open field, a novel object recognition test, and a T Maze task. Data from all 3 behaviors were used to perform hierarchical clustering analyses to delineate young average (black), aging unimpaired (AU; green), and aging impaired (AI; yellow) mice. Midbrains from these mice were dissected, and tissue underwent subcellular fractionation followed by proteomic analysis. **b)** PCA plots of all core matrisome proteins from membrane, cytoskeletal, and insoluble fractions. **c**) PCA plot of extracellular matrix proteoglycans from the membrane fraction (left). Average protein intensities of extracellular matrix proteoglycans in the membrane fraction separated by age and cognitive status (middle). Aging mice exhibited higher proteoglycan abundances (p < 0.001; n-Way ANOVA), and the aging impaired mice showed higher proteoglycan abundances compared to aging unimpaired mice (p < 0.05; post-hoc Tukey-Kramer). Right: heat plot of extracellular matrix proteoglycans abundances in the membrane fraction and bar plots of their relationship with cognitive status and age (* p < 0.05; linear probability model). **d**) PCA plot of synapse proteins from the membrane fraction (left). Average protein intensities of synapse proteins in the membrane fraction separated by age and cognitive status (middle). Right: heat plot of glutamate receptor abundances and bar plots of their relationship with cognitive status and age (* p < 0.05; linear probability model).

Principal component analysis (PCA) of entire proteomes from each fraction revealed the emergence of group separations between middle-aged impaired and unimpaired mice **(Ext. Data. Fig. 7),** suggesting links between the overall midbrain proteome and cognitive status of middle-aged mice. When restricting PCA analysis only to matrisome proteins, more prominent separations with respect to age and cognitive status were apparent across subcellular fractions (**Figure 6b**), supporting the idea that midbrain ECM status critically shapes cognition during aging. Among ECM subfamilies, proteoglycans showed more robust group separations in membrane and cytoskeletal fractions, while glycoproteins showed the greatest separation in insoluble fractions **(Ext. Data. Fig. 7)**. These observations suggest that both the abundance and subcellular localization/solubility of proteoglycans play roles in maintaining cognitive function with advanced age.

Proteoglycan abundance in membrane and cytoskeletal fractions was substantially higher in middle-aged mice compared to young (**Figure 6c; Ext. Data Fig. 7**). In membrane fractions, proteoglycan abundance was also significantly greater in middle-aged impaired mice compared to unimpaired (**Figure 6c**), whereas this separation in proteoglycan abundance with respect to cognitive status was not observed in either the cytoskeletal or insoluble fractions (**Ext. Data Fig. 7**). To evaluate what specific proteins drive these relationships, regression analyses between abundances of individual proteoglycans, age, and cognitive status were performed. This approach revealed that the predominant proteoglycans driving separations between middle-aged impaired and unimpaired mice were multiple HAPLNs (proteins that anchor proteoglycans to hyaluronan) and chondroitin sulfate proteoglycans (e.g., aggrecan, brevican, etc.; **Figure 6c**). Consistent with our prior proteomic analyses, many of these proteins increased in abundance during aging and showed significant positive correlations with age. Moreover, supporting the idea that this accumulation is not beneficial cognitively, many of these same proteins exhibited significant negative correlations with behavioral performance. As observed in our prior proteomic dataset, abundance of most synaptic proteins in these tissues did not change with age (**Figure 6d; Ext. Data. Fig. 8**), although multiple neurotransmitter receptors were negatively correlated with behavioral performance, indicating that middle-aged mice with less synaptic protein performed better on the cognitive battery (**Figure 6d; Ext. Data. Fig. 8**). This data provides an independent verification that having lower ECM proteoglycan and excitatory synapse abundances in the midbrain is cognitively beneficial for middle-aged mice and identifies the hyaluronan-proteoglycan matrix as a key modulator of cognitive aging phenotypes.

### Synapse and ECM abundance on dopamine neurons map onto cognitive aging phenotypes

Proteomic mapping in cognitively classified mice indicated that hyalectan abundances strongly align with cognitive phenotypes in middle-aged mice (**Figure 6**). In the subset of cognitively characterized mice (**Figure 6a**) that did not undergo proteomic analysis, we sought to independently validate this finding via immunohistochemistry for HAPLN1, an ECM link protein that directly interacts with hyaluronan, and aggrecan, an abundant chondroitin sulfate proteoglycan in the brain. HAPLN1-aggrecan complexes were found throughout the VTA, primarily near the surfaces of dopamine neurons and other neuronal or glial cells (**Figure 7a**), suggesting that HAPLN1 likely helps organize hyaluronan-proteoglycan interactions within the VTA. While field of view (FOV) HAPLN1 abundance did not differ across age, middle-aged impaired mice exhibited significantly higher HAPLN1 abundances compared to middle-aged unimpaired mice (**Figure 7b**). Aggrecan abundance did not differ with age or cognitive status (**Figure 7b**), although there were trends toward increased aggrecan abundance in middle-aged impaired mice. When analysis was restricted to zones within ∼0.5μm of dopamine neuron surfaces, we observed that HAPLN1 abundance was greater in middle-aged mice, and trending towards being significantly higher in impaired mice compared to unimpaired mice (**Figure 7c**). Aggrecan abundance on dopamine neurons did not differ with age or cognitive status. However, the abundance of aggrecan-HAPLN1 *complexes* on dopamine neuron surfaces was greater in middle-aged mice and also trending towards being significantly higher in impaired mice compared to unimpaired mice (p = 0.07; **Figure 7g; Ext. Data Figure 9**). These observations suggest that local ECM proteoglycan deposition on and around dopamine neurons critically informs overall cognitive performance during late middle age.

**Figure 7.**
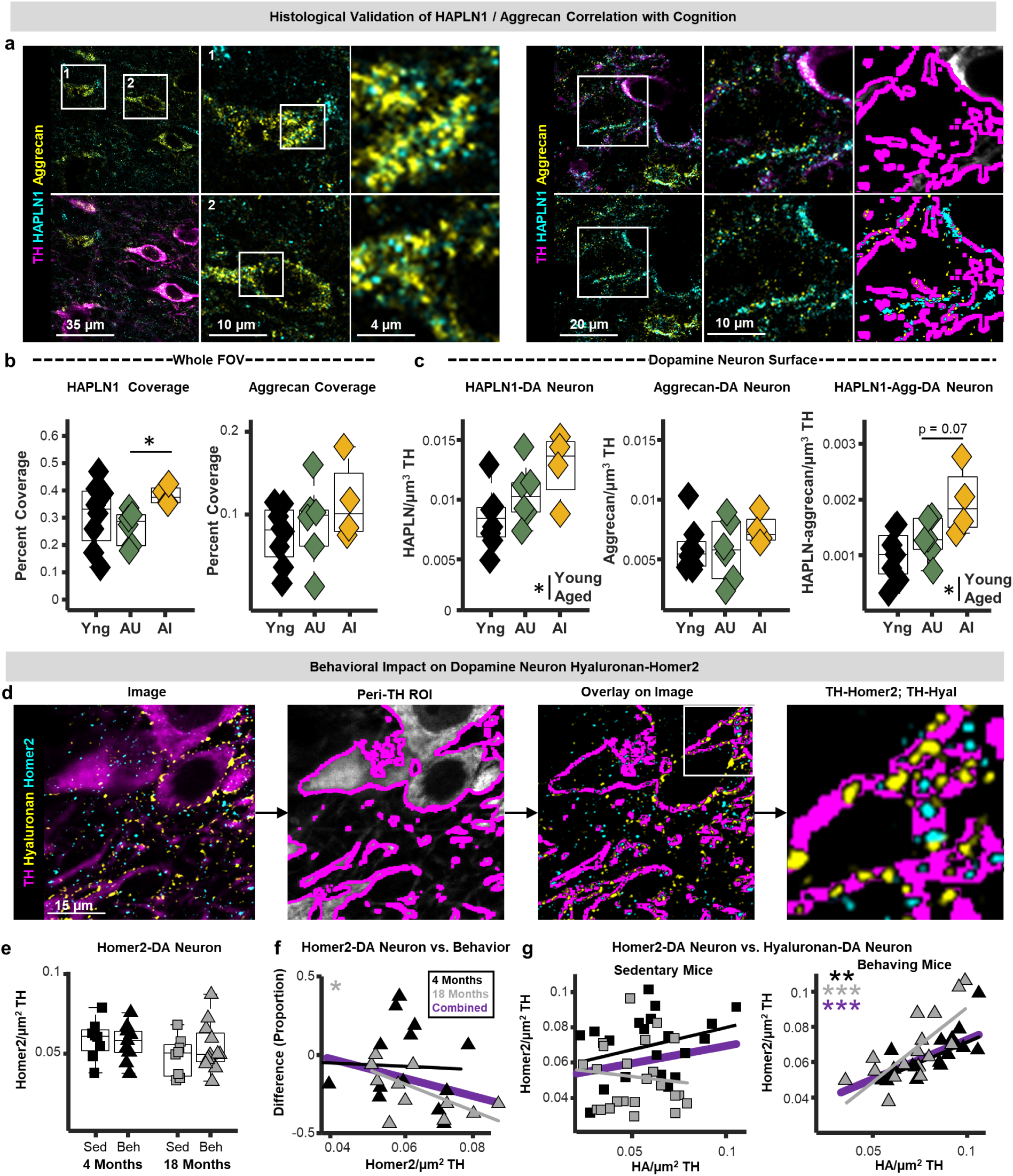
**a)** Left: example photomicrographs of immunohistochemically labelled hyaluronan and proteoglycan link protein 1 (HAPLN1), aggrecan, and tyrosine hydroxylase (TH) from the VTA of a late-middle aged mouse. Numbered images in the middle panel correspond to the numbered fields of view depicted by the white squares on the left. The right panels depict the fields of view depicted by the white squares in the middle panel. Right: example photomicrographs depicting HAPLN1 and aggrecan deposition on dopamine neuron surfaces, as well as the peri-dopamine neuron region of interest used to estimate the abundance of each protein at neuronal membranes in the VTA. **b)** Boxplots depicting field of view coverage of HAPLN1 and aggrecan in the VTA of young (black), aging unimpaired (AU; green), and aging impaired (AI; yellow) mice. AI mice exhibited significantly higher HAPLN1 coverage compared to AU mice (p < 0.05; n-way ANOVA with post-hoc t-test). **c)** Boxplots depicting HAPLN1-DA neuron, aggrecan-DA neuron, and HAPLN1-aggrecan-DA neuron puncta densities in the VTA of young, aging unimpaired (AU), and aging impaired (AI) mice. HAPLN1-DA neuron and HAPLN1-aggrecan-DA neuron densities increased with age (p < 0.05; 2-way ANOVA). AI mice tended to show higher HAPLN1-DA neuron and HAPLN1-aggrecan-DA neuron puncta densities compared to AU mice (p = 0.07; post-hoc Tukey-Kramer). **d)** Schematic of the analysis strategy used to examine hyaluronan and Homer2 on tyrosine-hydroxylase expressing dopamine neurons (Homer-DA neuron). **e)** Boxplots depicting homer-DA neuron puncta densities in young-adult (black) sedentary (squares) and behavior-trained (triangles) mice and late-middle-aged (grey) sedentary and behavior-trained mice. **f)** Relationship between homer-DA neuron puncta densities and performance on reward-based foraging task. A significant negative relationship was observed in middle-aged mice (p < 0.05; robust regression). **g)** Scatter plots of hyaluronan-DA neuron puncta densities plotted against Homer2-DA neuron puncta densities in sedentary and behavior-trained mice. No significant relationships were observed in sedentary mice, but significant positive relationships were seen in behavior-trained at both ages (young: p < 0.01; middle-aged: p < 0.01; all: p < 0.001 robust regression).

To further explore the role of dopamine-neuron-localized ECM in cognitive aging, we carried out similar analyses in tissue from mice trained in our 5-week food reward foraging paradigm (**Figure 5**), as well as sedentary controls (**Figure 7d**). Synapse abundance, as assessed via Homer2, on dopamine neurons did not differ between sedentary and behaving mice or across age. However, middle-aged animals with fewer dopamine-neuron-localized Homer2 puncta exhibited better task performance (**Figure 7e, 7f**), consistent with FOV analyses indicating that fewer VTA synapses correlated with better performance (**Figure 5k**). Hyaluronan abundance on dopamine neurons was lower in behavior-trained mice compared to sedentary mice (**Ext. Data Fig. 9**), agreeing with FOV findings that behavior training reduces hyaluronan density (**Figure 5g**). Importantly, both young and aging behavior-trained mice with more hyaluronan on dopamine neurons had significantly more Homer2 on dopamine neurons (**Figure 7g**). This correlation was completely absent in sedentary mice, suggesting that behavioral training engages hyaluronan-synapse remodeling around VTA dopamine neurons, and that failure to generate *and refine* these complexes is associated with cognitive decline in aging. Collectively, these histological observations identify hyaluronan and proteoglycan link proteins as promising targets for future mechanistic studies of cognitive resilience vs. decline.

### Microglia and ECM abundances independently correlate with cognitive aging phenotypes

Our findings in the context of microglial depletion (*Csfr1^ΔFIRE/ΔFIRE^*) and altered microglial aging trajectory (*Cx3cr1^EGFP/EGFP^*, Cx3cr1 KO) indicate that microglia may be poised to regulate abundance of VTA hyaluronan and proteoglycans (**Figure 4**). Moreover, in the context of normative aging, VTA microglial density was correlated with abundance of hyaluronan, and our results suggest that reduced microglial-ECM contact may contribute to ECM accumulations. To further probe potential links between microglial aging in mesolimbic circuits and cognition, we quantified VTA and NAc microglial densities in tissue from mice trained in the 5-week food reward foraging paradigm. As we showed previously, microglia densities were higher in the NAc compared to VTA and increased with aging only in the VTA (**Figure 8a, 8b**)^31^. Middle-aged mice with more VTA microglia exhibited worse task performance (**Figure 8c**), highlighting associations between this feature of microglial aging and cognition. Surprisingly, NAc microglia densities showed the opposite relationship, where mice with greater microglial densities exhibited better performance (**Figure 8c**). This suggests that regional microglial specializations and aging phenotypes uniquely impact the neuronal circuits in which they reside.

**Figure 8.**
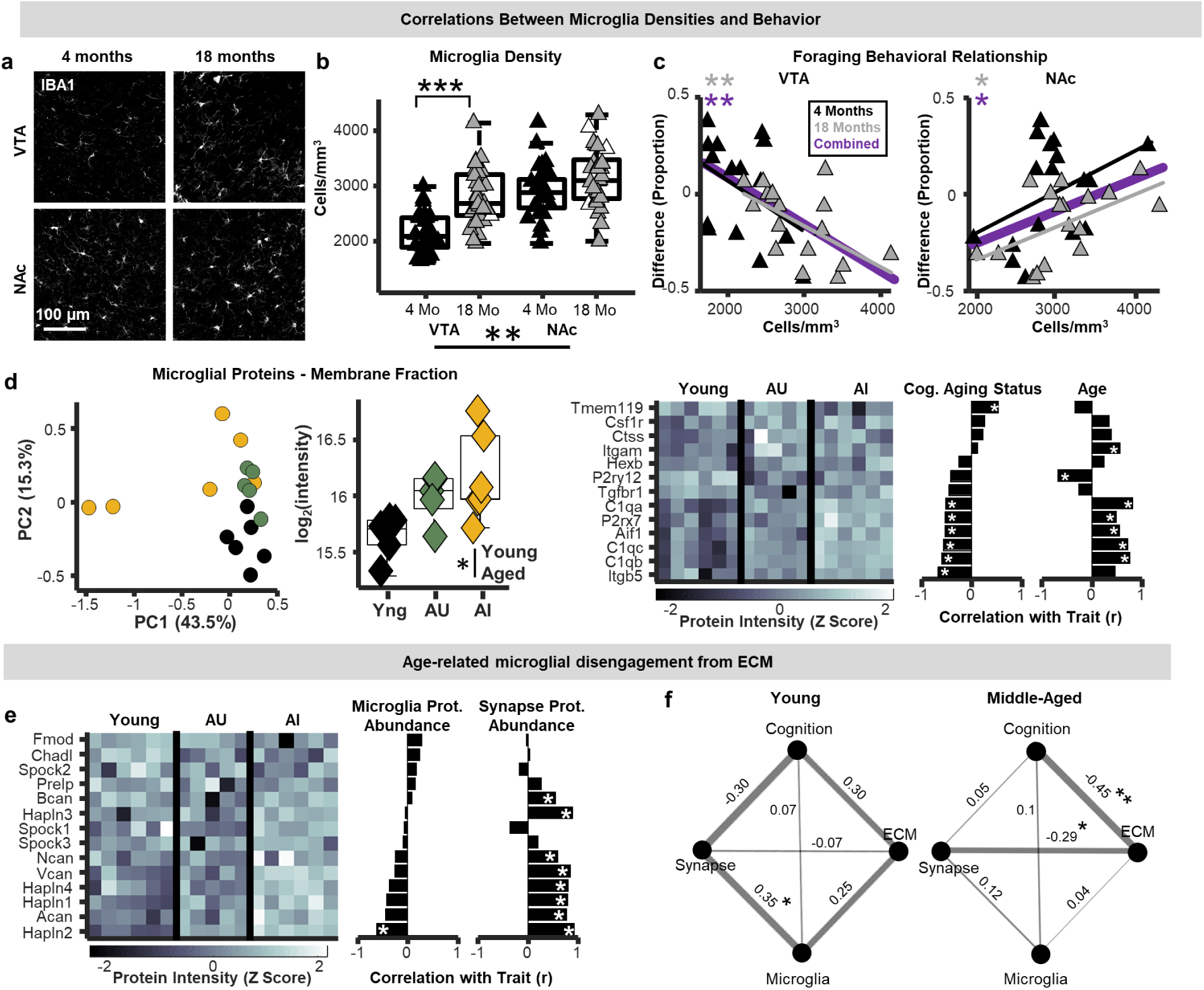
**a)** IBA1-positive microglia in the ventral tegmental area (VTA) and nucleus accumbens (NAc) of behavior-trained mice. **b)** VTA and NAc microglia densities in young-adult (black) and middle-aged (grey) mice. Greater microglia densities were observed in the NAc compared to VTA (p < 0.01; n-way ANOVA), and in the VTA, microglia densities were greater in middle-aged mice (p < 0.0001; post-hoc Tukey-Kramer). **c)** Relationship between VTA and NAc microglia densities and performance on reward-based foraging task. In the VTA, significant negative relationships were observed across all mice (purple; p < 0.01; robust regression) and in middle-aged mice (grey; p < 0.01; robust regression), whereas in the NAc, significant positive relationships were observed (all: p < 0.05; middle-aged: p < 0.05, robust regression). **d)** PCA plot of select microglia-enriched proteins from the membrane fraction (left). Average protein intensities of microglia-enriched proteins separated by age and cognitive status (middle). Microglia-enriched proteins were significantly more abundant in the aged midbrain compared to young (p < 0.05; n-way ANOVA). Heat plot of microglia-enriched protein abundances and bar plots of their relationship with cognitive status and age (* p < 0.05; linear probability model). **e)** Heat plot of ECM proteoglycan abundances from the membrane fraction and bar plots of their relationship with total microglia-enriched protein and synapse receptor abundances (* p < 0.05; linear probability model). **f)** Correlation network plots of relationships between cognition and ECM, synapse, and microglia abundances from both behavioral experiments in this study shown separately for young-aged and middle-aged mice. Numbers are r values and the width of connections between variables scales with the strength of their correlation (* p < 0.05; ** p < 0.01; linear regression).

Proteomic mapping also revealed increased abundance of multiple microglia-enriched proteins in the aged midbrain, and many of these increases were associated with age-related cognitive impairment (**Figure 8d**). The strong alignment between ECM abundance and cognitive aging phenotypes (**Figure 4**) prompts the question of whether microglia-ECM interactions influence cognitive aging, or whether their associations with cognition are independent of one another. To glean insights into this question, we first performed regression analyses between individual ECM proteoglycan abundances from cognitively-characterized aging mice (**Figure 6**), and total microglial and synapse receptor abundances in tissue proteomes from the same mice (**Figure 8e**). This analysis suggested that microglial protein abundances were not strongly aligned with most ECM proteoglycan abundances, although a significant negative correlation was observed with HAPLN2. Conversely, synapse receptor abundances showed strong positive relationships with HAPLN1-4 and all chondroitin-sulfate proteoglycans detected in this tissue (**Figure 8e**). To further evaluate these relationships, correlation network plots that integrate cognitive performance with microglial, ECM, and synapse abundances (measured via proteomics and histology) were created. In these plots, nodes represent individual features, and line thicknesses represent r values of pairwise correlations between traits. In young-adult mice, microglia abundances were strongly correlated to both ECM and synapse abundances, both of which showed relatively pronounced correlations with cognition (**Figure 8f**). In middle-aged mice, however, microglial relationships with both ECM and synapse abundances were drastically reduced, and ECM-synapse and ECM-cognition relationships were substantially higher (**Figure 8f**). Taken together, these analyses suggest that microglia and the ECM somewhat independently influence cognitive aging trajectories, and that ECM-synapse dynamics become more central to cognitive processing during normative aging.

## Discussion

As the predominant structure occupying the extracellular space, the ECM is positioned to play central roles in almost all neurological processes. Yet, brain ECM research remains nascent, and optimal strategies for observing and measuring the ECM’s complexity are still being defined. Revealing how the ECM impacts discrete brain structures – such as synapses – is a challenge for the field. ECM near synapses can regulate AMPA receptor diffusion, positioning of neuronal pentraxins^42,51^, extracellular ion concentrations, and access of phagocytic cells to synaptic elements^26,52^. Moreover, the “sweet spot” of optimal ECM abundance near synapses depends on context; appropriate ECM deposition may protect against synapse loss, but targeted ECM degradation is also essential for structural plasticity in support of learning and memory^7^. Advancing knowledge in this area is likely to reshape our understanding of synapse regulation in the aging brain and illuminate novel approaches to manipulate the brain ECM in support of healthy circuit function.

This report presents comprehensive proteomic mapping of the brain ECM during normative aging and reveals stark regional heterogeneity in ECM composition and age-associated remodeling across different basal ganglia nuclei. This argues that findings about the ECM in one brain region cannot be generalized to other brain regions and that similar ECM mapping of additional brain regions in a wide variety of contexts (CNS development, brain injury, brain cancer, neurodegeneration, etc.) is urgently needed. In general, basal ganglia matrisome protein abundance increased during aging, and histological examination of ECM proteoglycans and glycosaminoglycans confirmed this increase, aligning with previous biochemical and gene expression studies^53,54^. In the same tissue, we found no evidence of substantial aging-related synapse loss in the midbrain or striatum across multiple histological and proteomic datasets. This differs from previous reports that have shown age-related synapse loss in forebrain regions, including the prefrontal cortex and hippocampus^1,55,56^. These discrepancies could either reflect true brain region differences or that the present study primarily focused on middle-aged rather than geriatric mice. In terms of regional heterogeneity, the NAc had significantly more excitatory synapses and less ECM deposition compared to the VTA. This raises the possibility that NAc networks are more malleable, as the ECM is thought to restrict plasticity since, for example, the closing of developmental critical periods aligns with drastic increases in ECM abundance^57^. Furthermore, it may suggest that, although excitatory VTA synapse numbers remain stable with age, those synapses may also become increasingly rigid as VTA ECM deposition increases during aging.

Indeed, our data reveals multiple lines of evidence implicating the ECM as a key regulator of basal ganglia synapse function even in the absence of any changes to synapse number. For example, unbiased computational analyses of our proteomic data revealed that abundance of specific synaptic proteins strongly aligned with WGCNA modules containing ECM proteins, arguing that even in the absence of changes in synapse *number*, there is a critical role for the ECM in shaping synapse structure and composition. Histological analyses also revealed close anatomical proximity between postsynaptic markers (homer2) and hyaluronan, consistent with a structural and potentially functional relationship between ECM components and synaptic elements. Surprisingly, this analysis also revealed that synaptic protein abundances were not aligned with WGCNA modules containing complement proteins. Thus, while microglial synapse engulfment through complement tagging has been implicated in numerous disease contexts^23,58,59^, our data argues that local ECM status plays more prominent roles in regulating synapses during healthy brain aging. An important future direction for this research will be to examine relationships between basal ganglia ECM, immune, and synapse protein status using isolated synaptosomes, which allow for a more targeted quantification of synapse-associated proteomes^60^. Additionally, it will be critical to directly measure synaptic activity and capacity for synaptic plasticity and relate such measures to aging-related changes in ECM abundance and composition.

Our observation that middle-aged mice with *fewer* excitatory synapses exhibited better cognitive performance across both naturalistic foraging behaviors (paired with histology) and high-throughput behavioral assays (OF, NOR, T-maze - paired with proteomics) was rather unexpected. These observations challenge prevailing assumptions that more synapses automatically equates to better cognition^4,35^ and suggest that positive cognitive aging outcomes depend critically on the ability to refine and remodel synaptic networks. In support for this idea, we observed that repeated engagement in naturalistic foraging-based behavior had a net synaptogenic effect for both young and late-middle aged mice. However, mice that performed best on the foraging tasks had *fewer* excitatory postsynaptic puncta, implying that capacity to remodel rather than retain synapses is critical for optimal circuit function. Critically, in both behavioral experiments, greater synaptic protein levels and poorer cognitive performance were also associated with greater ECM abundance. These findings support a model in which excessive ECM accumulation may constrain the synapse remodeling and pruning required for optimal cognition^6,7,61,62^, and extends this framework into the context of normative brain aging. Moreover, these findings carry important implications for pathological aging contexts, including presymptomatic neurodegeneration or recovery from brain injury, where compensatory synaptogenic responses have been observed^63–65^. Our data argue that if these newly formed synapses are not appropriately refined, they may, at best, fail to facilitate appropriate circuit activity, and, at worst, exacerbate network dysfunction and cognitive decline through formation/retention of aberrant or inefficient synaptic connections. Nonetheless, an important limitation of our study is lack of information about the source and identity of synaptic inputs into the VTA and midbrain. Future studies leveraging connectivity mapping and *in vivo* functional imaging will be essential to fully understand how ECM-driven synaptic dynamics contribute to circuit-level adaptations and cognitive resilience in aging.

One unique feature of the VTA is that microglia within this region exhibit accelerated aging phenotypes, characterized by increases in proliferation and inflammatory factor production, compared to microglia in other basal ganglia nuclei^31^. Here, we provide critical replication of these VTA microglial aging patterns and build on this work by identifying what may be a previously unrecognized feature of midbrain microglial aging - a loss of capacity to contact and regulate the ECM. Several key findings support this hypothesis. First, while microglia densities and morphologies were significantly correlated with ECM deposition in young-adult mice, these relationships were lost by late-middle-age. Next, constitutive microglial depletion phenocopied normative age-associated increases in ECM abundance, arguing that microglia typically restrict ECM deposition and that this ability is lost during aging. Finally, perturbing microglial aging trajectories via deletion of Cx3Cr1 resulted in a failure to upregulate key ECM-regulatory genes that are upregulated in aging wild-type microglia^44^, resulting in greater ECM deposition compared to wild-type mice, and a loss of excitatory postsynaptic puncta by early middle age. These findings align with recent work showing that microglia regulate ECM structure to support synaptic plasticity in young-adult brains^7,25^, and argue that a loss of microglial regulation of the ECM during normative aging may contribute to excess ECM accumulation patterns we observe. An important future direction is to causally test how microglia modify ECM-synapse interactions in genetic mouse models with conditional manipulations to key microglial ECM-sensing proteins and ECM-degradative enzymes.

Our data also indicated several features of microglial aging that may impact cognition independently of microglial interactions with the ECM. For example, we found that abundance of complement proteins C1qA, C1qB, and C1qC was elevated with advanced age, consistent with previous work^67^. Via proteomic mapping of cognitive aging phenotypes, we also found that late-middle-aged mice with better cognitive performance had lower levels of complement proteins. Complement proteins could be influencing cognition via tagging synapses for phagocytic removal or via complement-ECM interactions that shape accessibility of synapses for pruning^68^. However, our proteomic analysis indicates that complement proteins do not strongly align with ECM- and synapse protein proteomic profiles during healthy aging. Moreover, recent findings indicate that C1q interacts with neuronal ribonucleoprotein complexes in an age-dependent manner, perturbing neuronal protein synthesis in a way that impacts cognition^69^. Together, these observations point to both ECM-dependent and ECM-independent molecular mechanisms by which microglia and immune molecules can potentially shape neural circuit function and cognitive resilience during aging.

This study lays a robust foundation for future research of glial-matrix biology in the context of cognitive aging. One of the more powerful elements of our study design was inclusion of two unique and sophisticated behavioral strategies that uniquely enabled us to link distinct cellular- and molecular-level observations about midbrain ECM and synapses to cognitive status of individual young and aging mice. For example, by including sedentary control mice in experiments where young-adult and middle-aged mice underwent reward-based behavioral training, we were able to detect behavior-induced hyaluronan matrix remodeling characterized by less ECM deposition around dopamine neurons that was aligned with synaptic phenotypes associated with better reward-based memory. This remodeling may reflect the engagement of somatodendritic dopamine release within the VTA in mice engaged in this task, as D1/D5 receptor activation has been linked to downstream protease release and ECM degradation^70^. More broadly, this observation demonstrates that ECM structure in the aging brain is not passively shaped by chronological age alone, but rather actively shaped by behavioral experience. This opens up exciting possibilities for non-invasive interventions (e.g., cognitive training or environmental enrichment)^71,72^ to modulate ECM states in support of cognitive function during aging.

Another critical finding for cognitive aging research was that protein expression patterns in tissue proteomes of late-middle-aged mice segregate solely based on unsupervised classification of their performance on canonical, high-throughput behavioral phenotyping tasks. This approach enabled us to directly link cognitive heterogeneity to underlying molecular states and identify the hyalectans as a specific family of ECM proteins strongly aligned with cognition in late-middle-aged mice. This behavioral classification strategy parallels work in the rat hippocampus linking histological and electrophysiological signatures of excitatory/inhibitory imbalance to poor cognitive aging^73^. More broadly, our work provides a scalable and unbiased experimental framework for appropriately sampling cognitive variability in aging mice in support of downstream experiments aimed at mechanistically dissecting cognitive resilience vs. decline.

While this work establishes novel pipelines for aligning ECM composition with cognitive performance, the immense complexity of the ECM leaves much to be uncovered. For example, our regional proteomic mapping experiments revealed aging-related changes in numerous perivascular ECM molecules (e.g., collagens and laminins) that have been linked to microglial activation patterns and remodeling of the neuronal ECM^74^. Given the important links between neurovascular health and cognitive function during aging^75,76^, it will be critical for future studies to focus on aging-related changes in the basement membrane-associated ECM in the context of cognitive aging. Moreover, our current methods provide limited insight into ECM assembly and spatial architecture beyond colocalization patterns. While we did not detect major shifts in solubility of most midbrain ECM proteins, we did observe aging-related differences in hyaluronan fragment size and filament length. Hyaluronan size has been shown to influence membrane excitability, diffusion properties in the extracellular space, and receptor signaling^26,77,78^. It will be important for future studies to understand how these subtler changes in ECM composition influence such physiological processes in the healthy aged brain. In this regard, there is evidence that unique post-translational modifications on ECM proteoglycans (i.e., hydroxylation, sulfation, and glycosylation) can alter key aspects of ECM physiology and its regulation of neuronal function during aging^10,11^. While our primary findings are based on relative comparisons within consistent experimental and analytical frameworks, our proteomic database searches did not include assessment of these ECM-relevant post-translational modifications, meaning that we likely underestimated matrisome protein abundance^79–81^ and cannot provide insights into more subtle changes in matrisome composition. A key future direction will be to implement proteomic pipelines and enrichment strategies that incorporate ECM-relevant post-translational modifications in order to capture the full complexity of the aging brain matrisome and its relationship with cognition.

Furthermore, there remains a pressing need for more precise and physiologically-relevant ECM manipulation strategies in neuroscience. Pharmacological approaches using compounds like 4-methylumbelliferone, a hyaluronan synthesis inhibitor, show inconsistent efficacy in the brain, require long treatment timelines, and provide limited temporal and spatial control^82,83^. Enzymatic ECM-degradation strategies have also been widely implemented in brain research; however, these treatments require invasive intracranial surgeries that induce mechanical injuries and the associated inflammatory and glial scarring responses^84,85^. This presents a major confound when examining normative aging phenotypes of neuron-extrinsic factors like the ECM and microglia^86,87^. These considerations underscore the need for new tools to manipulate specific ECM targets through viral approaches using retro orbital delivery^88^ or similar minimally invasive methods.

Finally, while this study identifies numerous aging-related ECM and microglial phenotypes associated with cognitive impairment, brain aging is not a passive process and there are numerous examples of adaptive aging-related neurobiological changes^48^. One potential example from the present study was that, while VTA microglial aging phenotypes aligned with worse behavioral performance, aging mice with more *NAc* microglia showed better goal-directed behavior. Hence, while some aspects of microglial aging represent vulnerabilities to circuit function, others may arise as adaptive responses that maintain neuronal network activity. As novel technologies emerge that allow for *in vivo* monitoring of specific ECM components, microglia-ECM interactions, and ECM-synapse interactions, such a framework (recognizing both detrimental and adaptive/beneficial aging-induced changes) will be critical in linking regional specializations in microglia-ECM-synapse dynamics with cognitive aging outcomes.

## Material and Methods

### Mice

This study uses C57Bl6 wild-type mice, CX3CR1^EGFP/+^ and CX3CR1^EGFP/EGFP^ mice, and Csf1r*^+/+^* and Csf1r^ΔFIRE/ΔFIRE^ mice^43^. C57Bl6 wild-type mice used for behavioral experiments and histochemical experiments were purchased from the National Institutes on Aging (NIA) colony (Bethesda, Maryland), and the mice used for regional proteomic mapping (**Figures 1-2**) were purchased from Jackson Laboratory (Bar Harbor, ME; stock #000664). CX3CR1^EGFP/EGFP^ breeders on a C57Bl6 background were originally purchased from the Jackson Laboratory (stock #005582) and crossed with C57Bl6 wild-type mice to obtain heterozygous (CX3CR1^EGFP/+^) mice and CX3CR1^EGFP/EGFP^ mice to obtain homozygous mice (CX3CR1^EGFP/EGFP^). In these mice, EGFP is knocked into the fractalkine receptor (CX3CR1) locus, which is a receptor expressed specifically by most myeloid-lineage cells, including microglia^90^. Previous work has demonstrated that EGFP expression in these mice is specific to microglial cells in the basal ganglia^18^. Csf1r^ΔFIRE/ΔFIRE^ breeders on a B6CBAF1/J background were originally purchased from Jackson Laboratory (stock # 032783) and crossed with C57BL/6 mice after which their offspring were interbred. These mice carry CRISPR/Cas9-generated deletion of the fms-intronic regulatory element (FIRE) of the Csf1r gene^43^.

For immunohistochemical and histochemical experiments in sedentary wild-type mice, up to 12 young-adult (4 months; 6 male, 6 female) and 12 late-middle aged mice (18 months; 6 male, 6 female) were used. Histochemical quantifications of synapse and ECM abundance in Cx3Cr1-deficient and knockout mice utilized 4 young-adult Cx3Cr1^EGFP/EGFP^ (3-4 months; 2 male, 2 female) and 4 middle-aged Cx3Cr1^EGFP/EGFP^ mice (12-15 months; 2 male, 2 female). Histological examinations comparing microglia-deficient mice with controls utilized 3 young-adult Csf1r^ΔFIRE/ΔFIRE^ mice (3-4 months, 2 females, 1 male) and 3 young-adult Csf1r^+/+^ mice (3-4 months, 1 female, 2 males). Quantitative proteomic experiments comparing ECM enrichment protocols utilized 8 young-adult wild-type mice (3-4 months). Midbrain tissue from 4 mice (2 male, 2 female) underwent chaotropic extraction and digestion protocol and tissue from 4 mice (2 male, 2 female) underwent the tissue fractionation protocol. Quantitative proteomic experiments of the midbrain and striatum of non-behaviorally characterized aging mice included 4 young-adult (3-4 months; 2 male, 2 female) and 4 aged (22-24 months; 2 male, 2 female) wild-type mice. Quantitative proteomic experiments of the midbrain of behaviorally-characterized mice included 6 young-adult mice (4 months; 3 male, 3 female), 5 middle-aged unimpaired (18 months; 2 male, 3 female), and 7 middle-aged impaired mice (18 months; 4 male, 3 female). Histological experiments of behaviorally characterized mice included 8 young-adult mice (4 months; 4 male, 4 female); 6 middle-aged unimpaired (18 months; 5 male, 1 female), 4 middle-aged impaired mice (18 months; 0 male, 4 female). Foraging-based behavioral testing and histological examinations utilized 18 young-adult (4 months; 9 male, 9 female) and 18 late-middle-aged (18 months; 9 male, 9 female) wild-type mice. Note that in all experiments measurements were taken from distinct samples from each animal. All mice within a given experiment were housed in the same vivarium with a normal light/dark cycle and were provided *ad libitum* access to food and water. Experiments adhered to protocols approved by the Animal Care and Use Committee and UCLA.

### Transcardial perfusion, immunohistochemistry, and histochemistry

Mice were deeply anesthetized in a covered beaker containing isofluorane and perfused transcardially with 1M phosphate buffered saline (PBS; pH 7.4) followed by ice-cold 4% paraformaldehyde (PFA) in 1M PBS. All perfusions were performed between 8:00 am and 12:00 pm to minimize the contribution that circadian changes may have on ECM, synapse, and microglial properties^91^. Brains were extracted immediately following perfusions and were allowed to post-fix for ∼4 hours in 4% PFA and then stored in 1M PBS with 0.1% sodium azide until tissue sectioning.

Coronal brain sections were prepared using a vibratome in chilled 1M PBS solution and stored in 1M PBS with 0.1% sodium azide until histochemical labelling. Sections containing the ventral tegmental area (VTA) and nucleus accumbens (NAc) were selected using well-defined anatomical landmarks at 3 distinct anterior-posterior locations per mouse. Free-floating brain sections were briefly rinsed in 1M PBS (5 minutes) and then permeabilized and blocked in a solution of 1% bovine serum albumin (BSA) and 0.1% saponin for 1 hour. For hyaluronan labelling, sections were then incubated with a biotinylated HABP lectin (1:250; Sigma-Aldrich cat: 385911) in the 1% BSA and 0.1% saponin block solution overnight at 4°C with mild agitation. Sections were washed in 1M PBS (4×10 minutes) and then incubated with streptavidin-conjugated AlexaFluor-647 in the 1% BSA and 0.1% saponin block solution for 2 hours. The tissue underwent another 4×10 minute wash in 1M PBS, and then was blocked again in 5% normal donkey serum with no additional permeabilization agent for 1 hour and was then incubated overnight with mild agitation in a solution containing different combinations of goat anti-GFP (1:1000; Frontier Institute cat: GFP-go-Af1480), rabbit-anti-IBA1 (1:500; Wako cat: 019-19741), chicken anti-TH (1:500; Aves cat: TYH), rabbit anti-Homer2 (1:2000; Synaptic Systems cat: 160 003), mouse anti-Vglut1 (1:2000; DSBH cat: N28/9), biotinylated WFA (1:100; Vector Labs, B-1355), rabbit anti-aggrecan (1:200; Sigma Aldrich AB1031), goat-anti HAPLN (1:200; R&D Systems cat: AF2608) and the biotinylated HABP lectin in 5% NDS solution. Sections were again washed in 1M PBS (4×10 minutes). Prior to secondary antibody incubation, all sections were treated with TrueBlack lipofuscin autofluorescence quencher (5%; Biotium cat: 23007) for 90 seconds followed by a 3×5 minute rinse in 1M PBS. Sections were then incubated in a secondary antibody solution containing combinations of chicken AlexaFluor-405 or chicken AlexaFluor-594, goat AlexaFluor-488 or goat AlexaFluor-647, mouse AlexaFluor-647, rabbit AlexaFluor-488 or rabbit AlexaFluor594, and streptavidin-conjugated AlexaFluor-647 in 5% NDS for 2 hours at room temperature with mild agitation. Experiments in which hyaluronan labelling was not performed were done in 1% BSA and 0.1% saponin throughout. The sections were again washed in 1M PBS (4×10 minutes), coverslipped (#1.5 thickness), and mounted using Aqua-Poly/Mount (Polysciences cat: 18606).

### Image acquisition and analysis

Fixed-tissue was imaged using a Zeiss LSM-700 confocal microscope using a 63x objective at a z interval of 0.3 µm or a Leica STELLARIS 5 confocal microscope using 20x (z = 1.5) and 63x objectives (z = 0.3). For quantification of hyaluronan and WFA fibril deposition patterns, 63x images were imported into Fiji image analysis software and underwent a background subtraction (rolling-ball radius = 15) and de-speckling. The processed images were then binarized using the ‘MaxEntropy’ setting and the proportion of a field of view covered by hyaluronan or WFA was calculated. Hyaluronan fibril densities and sizes were assessed using the 3D Object Counter plugin in Fiji using a minimum pixel cutoff of 10. Hyaluronan distribution regularity was quantified in the same images using the Fractal Dimension plugin in Fiji. HAPLN1 and aggrecan field of view coverage was calculated by importing images into Fiji image analysis software, applying a gamma correction and then a background subtraction (rolling-ball radius = 15). The processed images were then binarized using the ‘MaxEntropy’ setting and the proportion of a field of view covered by HAPLN1 or aggrecan was calculated. Densities of Homer2, VGlut1, putative Vglut1-Homer2 colocalization, putative hyaluronan-Homer2 interactions, and hyaluronan, HAPLN1, aggrecan, and Homer2 near TH-positive dopamine neurons were examined in 3-dimensions using custom-written MATLAB (Natick, MA) scripts using binarized images. Densities of ECM or synaptic components were normalized by the total surface area of TH signal within the field of view.

Microglia densities were calculated by manually counting microglial cell bodies within a z-stack using the Cell Counter plugin and then normalizing these counts by the volume of the image. Microglial morphological complexity was examined with 2-dimensional Sholl analysis using the SNT and Sholl Analysis plugin in Fiji^92^. Images containing IBA1 or GFP signal were first maximum intensity projected in the z dimension. A mark was placed in the center of each microglial soma and concentric radii were drawn from that point. The initial radius size was 7.5 µm and the step size of each radius after was 2.5 µm. The standard output values given by this plugin were used for analysis. Sholl intersection plots were made using MATLAB code. Manual reconstructions of microglia and the hyaluronan matrix were done by importing high-magnification (63x) confocal z-stack images into Imaris (Bitplane; Belfast, UK) software for reconstruction using the surfaces module. For each channel, a threshold that most accurately represented the signal was manually set and surfaces smaller than 1 µm^3^ were filtered out. Hyaluronan-microglia contacts were captured by filtering the hyaluronan surface using the GFP fluorescence histograms. The number of GFP-filtered hyaluronan aggregates was normalized by the total GFP within the field of view.

### Quantitative proteomics: sample preparation

Young and aged wild-type mice were anesthetized with isofluorane and perfused with 1M PBS. Brains were extracted and the midbrain and striatum were dissected using a scalpel, minced, triturated, washed in 1M PBS, and stored at -80°C until further processing. For solubility-based fractionation experiments, the tissue was processed using protocols from Naba et al., 2015^32^ that use a compartment protein extraction kit (Millipore cat: 2145). Triturated brain tissue was thawed on ice for ∼30 minutes before being homogenized in 2 ml/g of Buffer C (HEPES (pH 7.9), MgCl2, KCl, EDTA, Sucrose, Glycerol, Sodium OrthoVanadate) with protease inhibitors. This mixture was rotated on ice for 20 minutes and then spun at 20,000xg at 4°C for 20 minutes. The supernatant was removed and stored at -80°C until proteomic analysis. This fraction was considered the cytoplasmic fraction. The pellet was washed in 4 ml/g of the same buffer (HEPES (pH 7.9), MgCl2, KCl, EDTA, Sucrose, Glycerol, Sodium OrthoVanadate) with protease inhibitors, rotated for 5 minutes at 4°C, and spun at 20,000xg at 4°C for 20 minutes. The supernatant was removed and discarded. The pellet was then incubated in 1ml/g of Buffer N (HEPES (pH 7.9), MgCl2, NAcl, EDTA, Glycerol, Sodium OrthoVanadate), rotated at 4°C for 20 minutes, spun at 20,000xg for 20 minutes, and the supernatant was removed and stored at -80°C until proteomic analysis. This fraction was considered the nuclear fraction. The pellet was then washed in 1ml/g of Buffer M (HEPES (pH 7.9), MgCl2, KCl, EDTA, Sucrose, Glycerol, Sodium deoxycholate, NP-40, Sodium OrthoVanadate), rotated at 4°C for 20 minutes, spun at 20,000xg for 20 minutes, and the supernatant was removed and stored at -80°C until proteomic analysis. This fraction was considered the membrane fraction. The pellet was washed in 0.5 ml/g of buffer CS (Pipes (pH6.8), MgCl2, NAcl, EDTA, Sucrose, SDS, Sodium OrthoVanadate), rotated at 4°C for 20 minutes, spun at 20,000xg for 20 minutes, and the supernatant was removed and stored at -80°C until proteomic analysis. This fraction was considered the cytoskeletal fraction. The final remaining insoluble pellet was also stored at -80°C for proteomic analysis. This fraction was considered the insoluble fraction.

Chaotropic extraction and digestion experiments followed protocols from McCabe et al., 2023^33^, with the following modifications: Frozen samples were milled into powder in liquid nitrogen using a mortar and pestle. Approximately 5 mg of tissue was resuspended in a fresh prepared high-salt buffer (50 mM Tris-HCL, 3 M NaCl, 25 mM EDTA, 0.25% w/v CHAPS, pH 7.5) containing 1x protease inhibitor (Halt Protease Inhibitor, Thermo Scientific) at a concentration of 10 mg/ml. The samples were vortexed for 5 minutes and rotated at 4° C for 1 hour. The samples were then centrifuged for 20 minutes at 18,000xg at 4° C. The supernatants were discarded, and the pellets were resuspended in a high-salt buffer, vortexed for 5 minutes, and rotated again at 4° C for 1 hour. The samples were then recentrifuged for 15 minutes at 18,000xg at 4° C and the final pellets were used for proteomic analysis.

50-100ug protein was processed for mass-spectrometry analysis. Samples were resuspended in an equal volume of 100mM Tris-Cl (pH 8) and 8M urea. Proteins were then reduced with TCEP, alkylated using IAA, and cleaned using SP3 beads. Proteolytic digestion was performed overnight using lysC enzymes and trypsin. Peptides were subsequently cleaned using the SP3 protocol and eluted in 2% DMSO. Following elution, samples were dried in a speed vacuum and the resulting peptides were resuspended in 5% formic acid prior to liquid chromatography–tandem mass spectrometry (LC-MS/MS) analysis.

### LC-MS/MS parameters and database search

For quantitative proteomic experiments comparing ECM enrichment strategies (**Ext. Data Figure 1; Supplementary Table 1**) and experiments comparing midbrain and striatum proteomes of non-behaviorally characterized aging mice (**Figure 1; Supplementary Table 2**), LC-MS/MS was carried out as detailed in^93^. Approximately 200-500ng peptide amounts were subjected to LC-MS/MS analysis. Briefly, peptide separation was carried out using reversed-phase chromatography on a 75 μm inner diameter fritted fused silica capillary column, which was packed in-house to a length of 25 cm with 1.9 μm ReproSil-Pur C18-AQ beads (120 Å pore size). An increasing gradient of acetonitrile was delivered using a Dionex Ultimate 3000 nano-LC system (Thermo Scientific) at a constant flow rate of 200 nL/min. Tandem mass spectra (MS/MS) were acquired in data-dependent acquisition (DDA) mode on an Orbitrap Fusion Lumos Tribrid mass spectrometer (Thermo Fisher Scientific). Full MS1 scans were acquired at a resolution of 120,000, followed by MS2 scans at a resolution of 15,000. The raw LC-MS/MS data were analyzed using the MaxQuant computational platform^94^. The Andromeda search engine, integrated within MaxQuant, was employed for peptide identification against the UniProt reference proteome for *Mus musculus* (UP000000589). Search parameters allowed a maximum of two missed tryptic cleavages and included cysteine carbamidomethylation as a fixed modification and N-termination acetylation and methionine oxidation as variable modifications. A false discovery rate (FDR) threshold of 1% was applied at both peptide and protein levels. Label-free quantification (LFQ) was enabled, with a minimum LFQ ratio count set to one. Precursor ion and fragment ion mass tolerances were set at 20 ppm and 4.5 ppm, respectively. The resulting MaxQuant output files were subsequently used for statistical analysis to identify differentially enriched proteins. Raw data files have been deposited in the MassIVE proteomics repository under accession number MSV000096508. MS metrics and metadata information can be found in **Supplementary Table 3**.

For quantitative proteomic experiments of the midbrain of behaviorally-characterized mice (**Figure 6; Supplementary Table 4**), the cytosolic (CS), membrane (M), and insoluble (Insol) fractions were subjected to sequential reduction and alkylation steps using 5 mM tris(2-carboxyethyl)phosphine (TCEP) and 10 mM iodoacetamide, respectively. Following this, the protein aggregation capture (PAC) protocol, as described by Batth et al., (2019)^95^ was employed to purify the reduced and alkylated proteins. Proteins were then enzymatically digested overnight at 37 °C using Lys-C and trypsin proteases. The resulting peptide mixtures were dried completely and prepared for subsequent LC-MS/MS analysis. Dried tryptic peptides were resuspended in 5% formic acid and subjected to liquid chromatography-tandem mass spectrometry (LC-MS/MS) analysis using a Vanquish Neo ultra-high-performance liquid chromatography (UHPLC) system coupled to an Orbitrap Astral mass spectrometer (Thermo Fisher Scientific, Bremen, Germany). Briefly, peptide samples were introduced into a PepSep C18 reverse-phase analytical column (150 mm × 150 µm, 1.7 µm particle size), maintained at 59 °C, and separated using a trap-and-elute workflow. The UHPLC system employed mobile phase A consisting of water with 0.1% formic acid and mobile phase B consisting of acetonitrile with 0.1% formic acid. Peptide separation was achieved using a 15-minute chromatographic gradient with the following composition: 5% B from 0–1 min at a flow rate of 2.45 µL/min; a linear increase from 5% to 15% B from 1–5 min at 1.75 µL/min; 15% to 25% B from 5–12.6 min at 1.75 µL/min; 25% to 38% B from 12.6–13.6 min at 1.75 µL/min; and a rapid gradient from 38% to 80% B between 13.6–13.7 min at 2.45 µL/min, followed by a hold at 80% B until 15 min, maintaining the flow at 2.45 µL/min.

Mass spectrometric acquisition was carried out in data-independent acquisition (DIA) mode on the Orbitrap Astral instrument operating in positive electrospray ionization mode. MS1 survey scans were acquired across an m/z range of 380–980 with a resolution of 240,000, using a normalized AGC (automatic gain control) target of 500% and a maximum injection time of 3 ms. For DIA, sequential isolation windows of 4 m/z were used to comprehensively cover the 380–980 m/z range. Fragment ion (MS2) spectra were acquired at a resolution of 80,000, with a normalized higher-energy collisional dissociation (HCD) energy of 25%, an AGC target of 500%, and a maximum injection time of 7 ms. The raw Thermo .RAW files were analyzed using DIA-NN software, searching against an in silico predicted spectral library generated from the Mus musculus reference proteome (UniProt ID: UP000000589), as described by Demichev et al., (2020)^96^. The resulting DIA-NN outputs were processed using FragPipe Analyst to identify differentially expressed proteins, following the workflow outlined by Hsiao et al., (2024)^97^. All raw data files have been deposited in the MassIVE proteomics repository under accession number MSV000096508. MS metrics and metadata information for these data can be found in **Supplementary Table 5**.

### Quantitative proteomics: analysis

Protein intensity values for all samples were batch corrected via quantile normalization and median centering. For regional mapping experiments of ECM protein abundance across subcellular/solubility fractions (**Figure 1c-1e**), age-related shifts in protein solubility were examined by calculating the fold-change of individual ECM protein abundance (log2-normalized protein intensities) that were detected across at least 2 distinct fractions (fold change relative to the more insoluble fraction), and comparing these fold-changes between young-adult and aged mice. For regional mapping experiments of total ECM protein abundance, and WGCNA analysis (see below), data for each identified protein was summed across solubility fractions for each mouse and log2 normalized for analysis to estimate total protein intensity. For proteomic mapping of cognitive phenotypes, log2-normalized protein intensities detected in membrane, cytoskeletal, and insoluble fractions were examined independently. This proteomic mapping was only performed on midbrain tissue. To be included in the analysis, a protein needed to be detected in 75% or more of the samples within a brain region. Fold changes were calculated as the ratio of the aged signal relative to the young signal. Statistical significance was assessed using unpaired t-tests with an alpha-level of 0.05. Annotations for different protein classes were downloaded from publicly available online databases. ECM protein annotations were downloaded from the Matrisome project database^98^, innate immune system protein annotations from the InnateDB database^99^, and synapse protein annotations from the SynGo database^100^.

### Quantitative proteomics: weighted gene correlation network analysis (WGCNA)

To examine relationships between age, cognitive status, synaptic, matrisome, and innate immune proteins, proteomic data underwent unsupervised clustering using Weighted Gene Correlation Network Analysis (WGCNA) in R^34^. For the regional mapping experiment, processed log2(intensity) data from the midbrain and striatum were used for network construction. One striatal sample from an aged female mouse was identified as an outlier and removed from this analysis. WGCNA provides modules of covarying proteins that are agnostic to traits of the samples (i.e., age, region, sex). The distribution of synaptic, matrisome, and immune proteins across the protein modules was used to identify modules of interest for further analysis. Relationships between modules and external traits or protein abundance were examined using the module-trait relationship code provided in the WGCNA R package. Process enrichment analysis of the protein modules was conducted in Metascape^101^ using default parameters. For the proteomic mapping of cognitive phenotypes experiment, only proteomic data from the midbrain was used for WGCNA analysis. Using age and cognitive status (see below) as traits, protein-trait relationships were derived using the module-trait relationship code provided in the WGCNA R package.

### Mining of published RNAseq data from wild-type and Cx3Cr1 deficient microglia

Publicly available bulk RNA-seq datasets of isolated microglia from Cx3cr1 knockout (Cx3cr1⁻/⁻), Cx3Cr1 heterozygous (Cx3cr1^+^/⁻), and wild-type mice were retrieved from Gyoneva et al., 2019^44^. Analysis was restricted to a curated list of matrisome genes, defined according to the MatrisomeDB^98^, encompassing core ECM proteins and ECM regulators. Fold changes of aging-induced changes in matrisome genes were extracted from datasets comparing 2 month and 12 month old Cx3Cr1-knockout and wild-type mice. Additionally, fold changes of matrisome gene expression between Cx3Cr1 heterozygous and Cx3Cr1 homozygous knockout (Cx3cr1⁻/⁻) mice (2 months old) were examined. This targeted approach enabled the identification of Cx3cr1-dependent alterations in microglial ECM-associated gene expression.

### Foraging-based behavioral testing procedures

For the foraging-based behavioral experiment, behavioral testing was conducted in a large square open field (120 cm x 120 cm) with 35 cm high walls made from transparent Plexiglas. On one wall, there was a 20 cm x 20 cm opening where a start box equipped with a guillotine door was placed. This start location did not change across testing sessions. Mouse bedding was placed on the floor of the arena, and 2 intra-arena reference cues were placed at two fixed locations for the entire experiment. To also provide fixed distal landmarks to the mice, black corrugated plastic was placed on two adjacent walls in the arena, and white corrugated plastic was placed at the other two. Plexiglas dishes were filled with Sandtastik play sand (Sandtastik Products LLC, Port Colborne Ontario) and were used to plug 5TUL purified rodent tablets into (TestDiet; cat: 1811142; Quakertown, PA) to engage foraging behavior. Identical feeders without reward pellets were used as distractor feeders to test goal-directed memory.

The feeders were placed at 1 of 4 locations within the arena. All feeders were sham baited with powdered reward pellets (5% by weight). Mouse behavior was recorded using a 1.3 megapixel, low-illumination overhead camera (ELP; Shenzhen, Guangdong, China) and Bioserve Viewer software (Behavioral Instruments; Hillsborough, NJ).

Young-adult (3-4 months) and late-middle-aged (16-18 months) mice were ordered in 3 cohorts of 20. Twelve mice from each cohort then underwent behavioral testing and the other 8 lived with the behavior-trained mice but did not undergo testing (sedentary controls). Thus, in total this experiment used 24 sedentary mice (12 young, 12 middle-aged; 6 male and 6 female in each group) and 36 behavior-trained mice (18 young, 18 middle-aged; 9 male and 9 female in each group). Sample sizes were determined based on variability in pilot data and from published data on a similar task from which this one was derived^102^.

Behavior-trained mice were handled daily for 1 week prior to testing. During week 1 of testing (habituation phase) mice were placed within the start box for ∼2-3 minutes prior to the guillotine door opening to allow entry into the arena. A single sand well with 4 reward pellets was placed at one of the reward locations and the mice were allowed 5 minutes to explore the arena and retrieve the reward. This procedure was repeated daily for 5 sessions for each mouse and the location of the rewarded feeder was changed daily and was counterbalanced within a day across mice. During weeks 2 and 3 (training phase), the mice again entered the arena via the start box and foraged for reward in a single sand well at one location in the arena for 5 minutes (encoding). Following a 30-minute delay period, mice reentered the arena with the rewarded feeder in the same location and 3 unrewarded feeders in the other locations for 2 minutes (retrieval). Note that this is a win-stay strategy. Again, the location of the rewarded feeder was changed daily for each mouse. Each mouse underwent 10 training sessions across 2 weeks of testing. In the final 2 weeks of testing (test phase), mice repeated the encoding procedures described above. Following either 2- or 24-hour delays, mice underwent a probe test in which 4 unrewarded feeders were placed within the arena, 1 at the previously rewarded location. After 2 minutes the mice exited the arena through the guillotine door and the experimenter placed reward pellets in the previously rewarded location. The mice then reentered the arena and consumed the food reward to minimize extinction. Again, rewarded locations changed across sessions, and the 2- and 24-hour delays were interleaved to eliminate practice effects between the two conditions.

### Foraging-based behavioral testing data analysis

Mouse behavioral tracking was collected using Bioserve Viewer software in 1-second timestamps and was used to determine the time of entry and exit to various zones of interest within the arena. These zones included the reward feeder locations, and a box that separated the arena’s perimeter from the center. Data were exported from the Bioserve Viewer software and loaded into Matlab for behavioral analysis using custom-written scripts. From this data the following output measures were derived: average running speed, start-box exit latency, proportion of time on the arena’s perimeter, number of feeder visits, proportion of feeder visits to the correct feeder, the relative proportion of time in correct and incorrect feeders, and the number of errors prior to correct feeder visits. Note that because there were 4 foraging locations, chance performance during probe trials is 25% for proportional measures and 1.5 for error measures. Probe trial data were used as the estimate of goal-directed memory function in these mice.

The temporal progression of the emergence of foraging behavior was tracked across sessions for each animal with a state-space modeling approach using Bernoulli observation models^103^. Mice were considered to have foraged if they consumed at least 1 of the 4 treats within the arena during the encoding sessions. Using this binary assessment (1 = consumed, 0 = non consumed), a learning curve, its 90% confidence bounds and the trial at which consistent foraging was observed were determined. This trial was defined as the point in which the lower bound of the 90% confidence interval exceeded chance (50%) and stayed above chance for the remainder of the experiment.

### Behavioral battery for proteomic mapping of cognitive phenotypes

For behavioral characterization experiments using spontaneous rodent behaviors, 12 young-adult (4 months; 6 male and 6 female) and 24 middle-aged (18 months; 12 male and 12 female) were used. First, all mice underwent an open field test to assess general locomotor activity and anxiety-like behavior using a square arena (60cm x 60 cm with opaque walls 30 cm high). Mice were placed in the center of the arena and allowed to explore freely for 10 minutes. Their behavior was tracked with an overhead camera using Bioserve Viewer software, and the floor of the arena was digitally divided into central and peripheral zone to calculate the proportion of time spent in the center and periphery of the arena. The arena was cleaned with 70% ethanol and water between mice. Following open-field testing, all mice then underwent a novel object recognition (NOR) task in the same behavioral apparatus. Because testing was done in the same apparatus, the open field test also served as habituation for the NOR test. Thus, NOR testing began with familiarization of two identical objects (either two LEGO stacks or two PVC pipes of equal height and width, counterbalanced across subjects) that were placed symmetrically near the corners of the arena. Mice again were allowed to explore for 10 minutes. Using a separate group of aging mice, we confirmed that mice did not inherently prefer one object over the other by placing one of each object within the arena and allowing mice to explore for 10 minutes. After a retention interval of 2 hours, NOR mice underwent the test phase of the task where one familiar object was replaced with a novel object. The position of the novel object (left/right) was counterbalanced across subjects to control for side preference. Again mice explored for 10 minutes. Object exploration was defined as the mouse directing its nose, within 2 cm from the object. Using this data, a discrimination index was calculated as follows: (time with novel – time with familiar) / (total time with both objects). Objects were cleaned with 70% ethanol and water between trials. Following NOR testing, mice underwent a T-maze spontaneous alternation test. This testing was done in a t-shaped maze made of white corrugated plastic comprising of a start arm and two goal arms. Each arm was 30 cm long, 10 cm wide, and 25 cm tall. During the first trial, mice were forced into either a right or a left decision using a removable barrier. This forced choice was counterbalanced between mice. During all subsequent trials, mice were allowed to freely choose between the two goal arms. After entering one of the goal arms, mice were returned to the start arm for the subsequent trial. The procedure was repeated for 10 consecutive trials, with an inter-trial interval of ∼30 seconds. An alternation was recorded when the mouse chose the goal arm opposite to the one it entered on the previous trial, and an alternation index was calculated as follows: number of alternations / total number of trials.

### Classification of middle-aged unimpaired and impaired mice

To behaviorally classify middle-aged mice as cognitively impaired or unimpaired, behavioral data from open field, NOR, and the T-maze were used for unsupervised hierarchical clustering analysis. For each mouse, a behavioral performance vector was created using the proportion of time on the arena perimeter from the open field test, the discrimination index from the NOR test, and the alternation index from the T-maze. All data was z-scored prior to constructing these vectors to ensure equal weighting across features. Hierarchical clustering was performed in MATLAB using the ‘linkage’ function with Ward’s method (minimizing total within-cluster variance) and Euclidean distance as the similarity metric. A dendrogram was generated using the ‘dendrogram’ function to visualize hierarchical relationships between subjects. Clusters were identified by applying a threshold to the dendrogram (via cluster function with k=2), resulting in two distinct clusters. Middle-aged mice that fell within the cluster with the majority of young mice (75%) were considered unimpaired and the middle-aged mice that fell in the other cluster were considered impaired. This data-driven behavioral classification was then used to examine group-level differences in proteomic and histological data.

### Statistical analysis

Statistical analyses were done using either MATLAB or R. Statistical significance of proteomic protein-intensity data was evaluated using unpaired t-tests. Immunohistochemical data was statistically analyzed using multi-way ANOVAs with Tukey’s post-hoc tests where applicable, and foraging behavior data was assessed using repeated-measures ANOVAs and unpaired t-tests. All statistical tests included age and sex (wild-type) or age, sex, and genotypes (Cx3Cr1 KO) as covariates and were two-sided. Relationships between anatomical variables and behavioral variables, or between different anatomical variables were assessed using robust regression analyses. Relationships between proteomic data and cognitive status were examined using point-serial correlation analysis (regression with one binary variable). To visualize relationships between all behavioral, anatomical, and proteomic variables created in both behavioral experiments, a correlation-based network analysis was performed in MATLAB. Here, a pairwise Pearson correlation matrix was computed across all behavioral measures using the ‘corrcoef’ function with pairwise deletion for missing data. Self-correlations on the diagonal (r = 1) were set to zero to exclude self-links in the resulting network. This network was then visualized as an undirected graph using the ‘graph’ function. For all experiments, a significance cutoff of p < 0.05 was used. Potential sex differences in ECM, microglia, and excitatory synapse status were evaluated in all cases, and all statistical parameters and test statistics are available in **Supplementary Table 6**.

## Data Availability

Statistical source data is provided with this paper in supplementary files. Raw quantitative proteomics datasets were deposited in the MassIVE online database (accession number MSV000096508). Any other data is available upon request from the authors.

## Code Availability

Custom code used for analyses are available on CodeOcean (capsule number: 18-559067-0).

## Extended Data Figures and Figure Legends

**Extended Data Figure 1.**
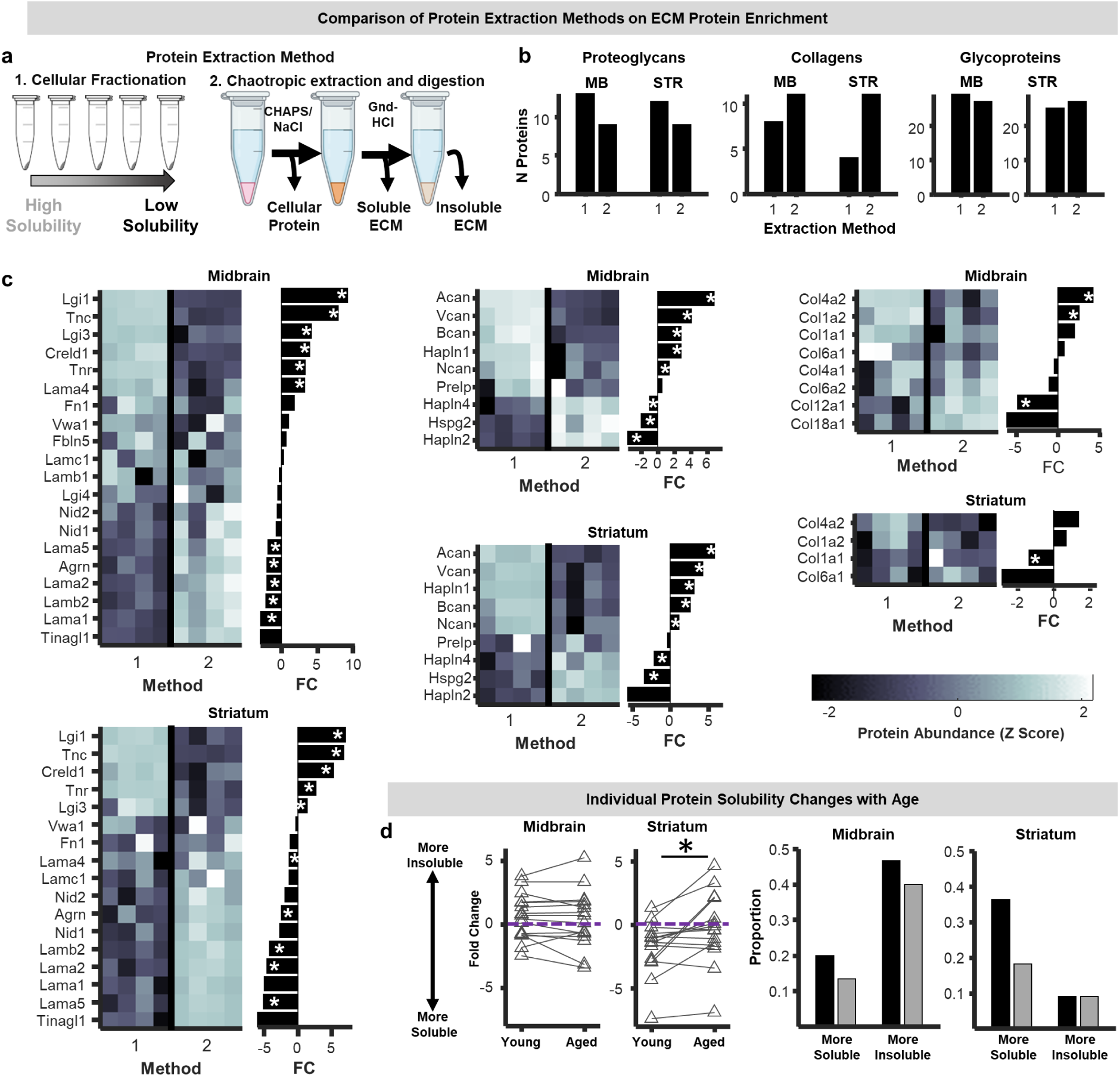
**a)** To compare distinct ECM protein enrichment strategies, the midbrain and striatum of young-adult (3 months) mice were dissected and underwent either 1: solubility-based subcellular fractionation (n=4) or 2: chaotropic extraction and digestion (n = 4) or protocols. **b)** Comparison of the number of core matrisome proteins detected using each approach. Extraction method 1 refers to solubility-based fractionation and extraction method 2 refers to chaotropic extraction and digestion **c)** Heat maps of protein abundances (z scored) and bar plots of corresponding fold-changes with age for ECM glycoproteins, proteoglycans, collagens, and ECM regulators detected using the two approaches. (* p < 0.05 - unpaired t-test). **d)** Left: Paired plots of fold changes of individual matrisome proteins detected in the insoluble fraction that were detected in at least one of the subcellular/soluble fractions using the fractionation approach for young and aged mice. Fold-changes were calculated with respect to the insoluble fraction. There was a significant shift towards greater insolubility in the aged striatum (p < 0.05; paired t-test). Purple lines denote a fold change of 0, indicating no difference in abundance between insoluble and soluble fractions. Right: bar plot showing the proportion of matrisome proteins showing differences in solubility (upregulated and downregulated shown separately) in the young and aged mice.

**Extended Data Figure 2.**
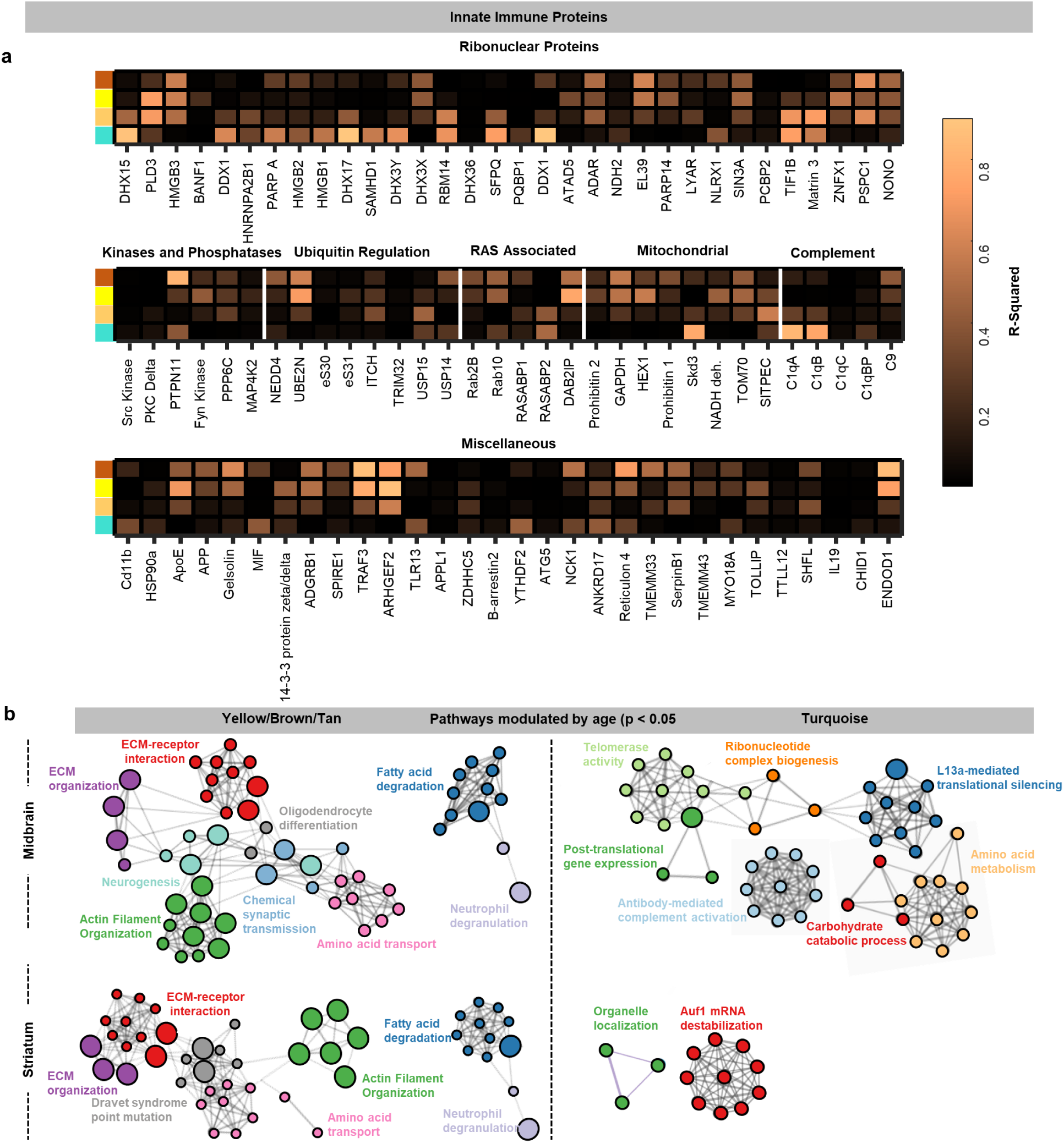
**a)** Heatmaps representing correlations (r-squared values) between brown, yellow, tan, and turquoise module eigengenes and abundances of specific innate immune proteins separated by functional classification (ribonuclear proteins, kinases and phosphatases, ubiquitin-associated proteins, mitochondrial proteins, complement proteins, and other miscellaneous innate immune proteins). **b)** Network plots of process and pathway enrichment terms associated with all tan/yellow/brown module (left) and turquoise module (right) proteins whose abundances were modulated by age (p < 0.05) in the midbrain and striatum.

**Extended Data Figure 3.**
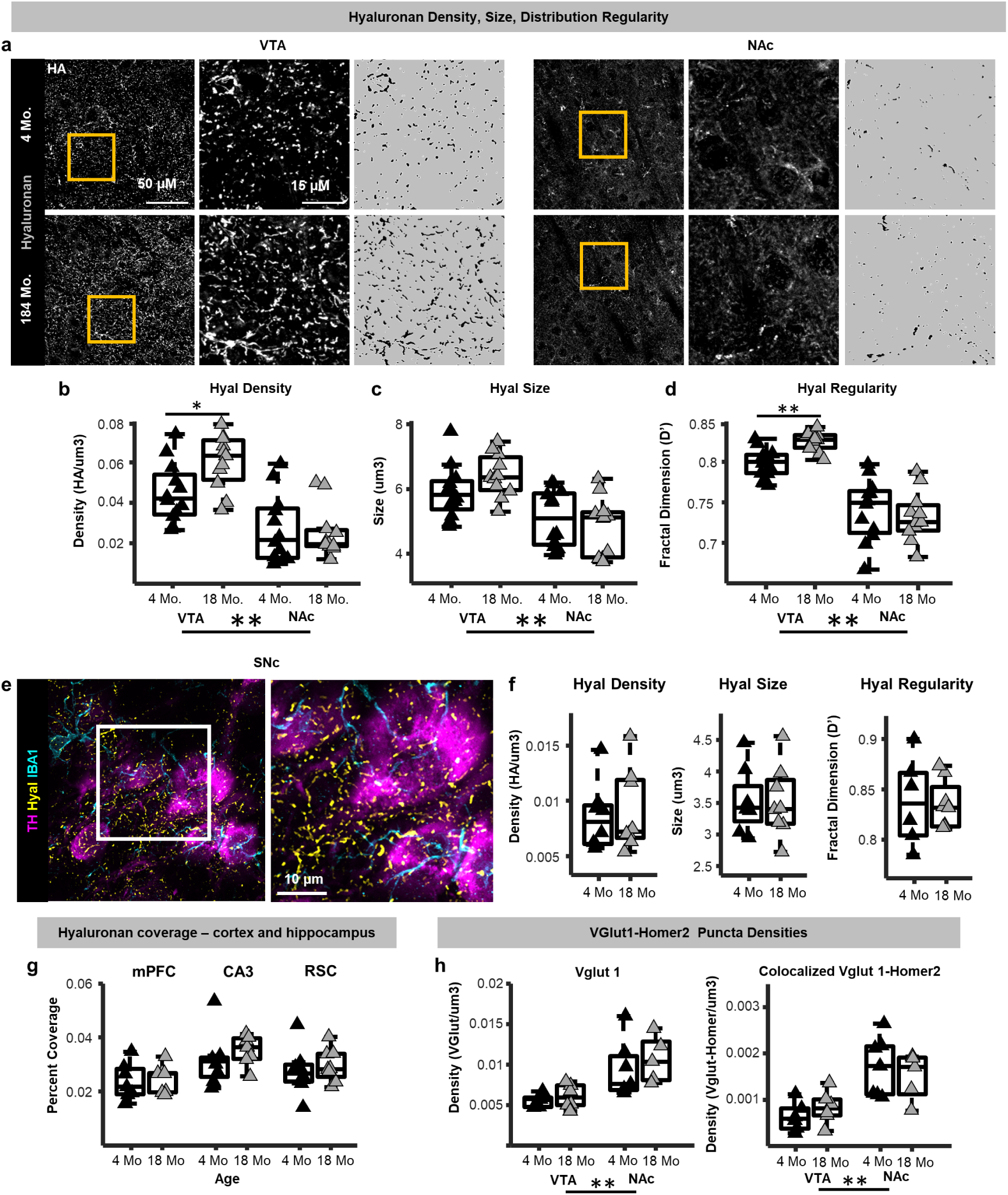
**a)** The ECM scaffold hyaluronan (Hyal) was analyzed in two critical regions of the mesolimbic dopamine system, the ventral tegmental area (VTA) and nucleus accumbens (NAc). **b)** Example photomicrographs of histochemically labelled hyaluronan in the VTA and NAc of a young-adult (4 months) and late-middle-aged (18 months) mouse (left panels). Middle panels show zoomed in images of the areas within the yellow squares in the left panels. The right panels depict binarized images of the hyaluronan matrix used for quantification. **c-e)** Boxplots depicting **c)** hyaluronan fibril densities, **d)** median hyaluronan fibril sizes, and **e)** hyaluronan distribution regularity (Fractal Dimension plugin – Fiji) in the VTA and NAc of young-adult (black) and late-middle-aged (grey) mice. Higher fractal dimension values (D’) indicate more spatial regularity of the hyaluronan matrix. Hyaluronan fibril density, median size, and distribution regularity were greater in the VTA compared to NAc (Density: p < 0.001; Size: p < 0.001; Regularity: p < 0.001; n-way ANOVA). Late-middle-aged mice exhibited significantly greater VTA hyaluronan fibril density (p < 0.05; post hoc t-test) and distribution regularity (p < 0.01; post-hoc t-test). **e)** Example photomicrograpghs of histochemically labelled hyaluronan, tyrosine hydroxylase (TH)-positive dopaminergic neurons, and IBA1-positive microglia in the substantia nigra pars compacta (SNc). **f**) SNc hyaluronan fibril densities, sizes, and distribution regularity were not different between young-adult (4 months) and late-middle-aged mice (18 months). **g)** Hyaluronan fibril field of view coverage in the mPFC, CA3 region of the hippocampus, and retrosplenial cortex (RSC) of young-adult (4 months) and late-middle-aged mice (18 months). **h)** Left: densities of VGlut1 puncta in the VTA and NAc of young-adult and middle-aged mice. There was a greater density of VGlut1 in the NAc compared to VTA (p < 0.01; n-way ANOVA). Right: densities of VGlut1-homer2 colocalized puncta in the VTA and NAc of young-adult and middle-aged mice. There was a greater density of VGlut1-homer2 colocalized puncta in the NAc compared to VTA (p < 0.01; n-way ANOVA).

**Extended Data Figure 4.**
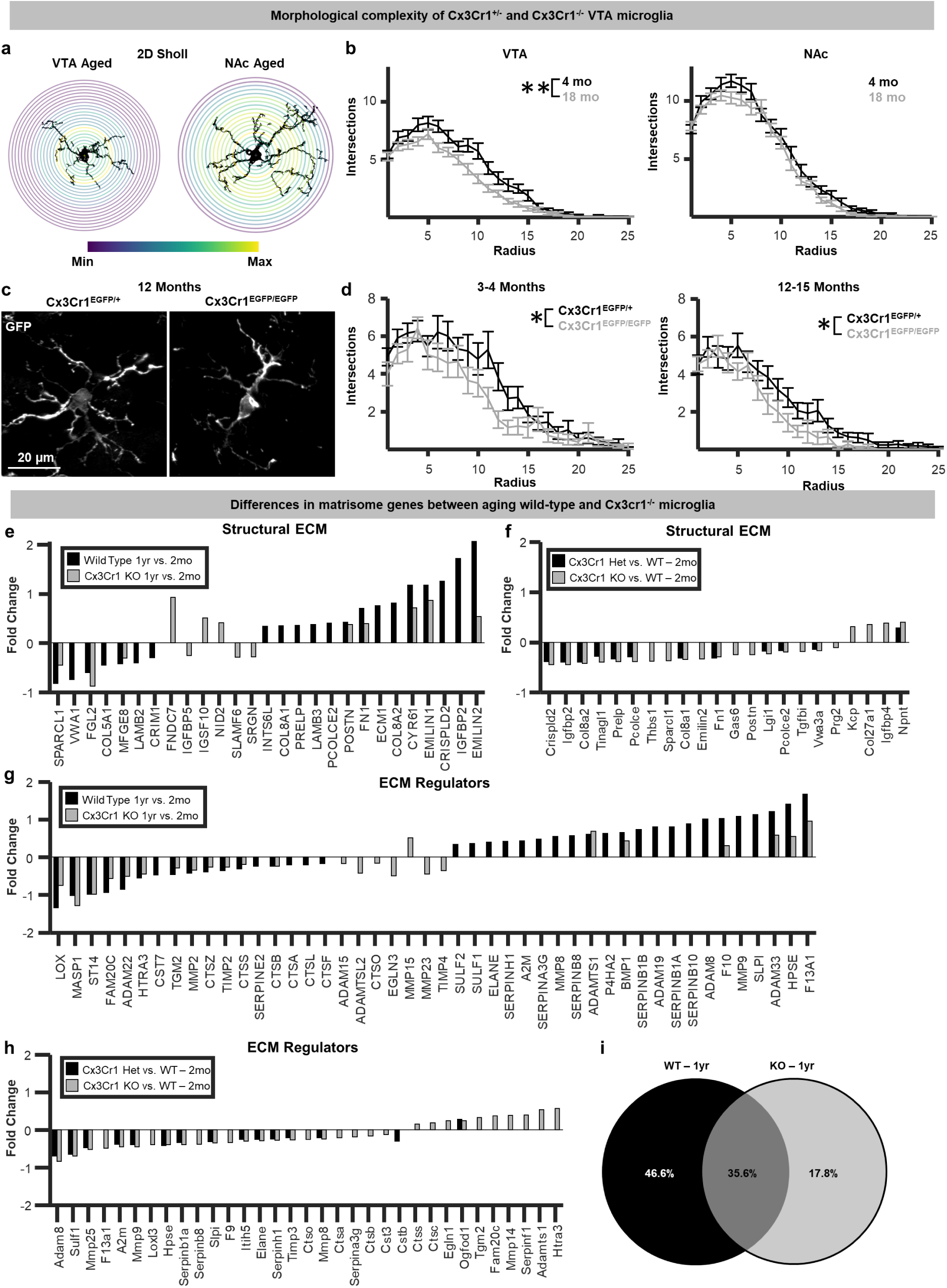
**a)** 2-dimensional Sholl analysis was performed on ventral tegmental area (VTA) and nucleus accumbens (NAc) IBA1-positive microglia in young-adult (4 months) and late-middle-aged (18 months) mice. **b)** Sholl intersection plots for VTA and NAc microglia for young-adult and late-middle-aged mice. Middle-aged VTA microglia exhibited a reduced morphological complexity compared to young adults (p < 0.01; n-way ANOVA). **c)** Example photomicrographs of GFP-positive microglia in the VTA of 12-month-old Cx3Cr1^EGFP/+^(het) and Cx3Cr1^EGFP/EGFP^ (KO) mice. **d)** Sholl intersection plots for VTA microglia from young-adult (3-4 months) and early-middle-aged Cx3Cr1-het and Cx3Cr1-KO mice. Cx3Cr1 knockout microglia exhibited a reduced morphological complexity compared to both young-adult and middle-aged mice (p < 0.01; n-way ANOVA). **e)** Fold changes of structural matrisome genes that were significantly differentially expressed between 1-year and 2-month wild-type (black) and Cx3Cr1-knockout (grey) microglia (from Gyoneva et al., 2019). **f)** Fold changes of structural matrisome genes that were significantly differentially expressed between 2-month Cx3Cr1 heterozygous (black) and Cx3Cr1-knockout (grey) microglia. **g)** Fold changes of regulatory matrisome genes that were significantly differentially expressed between 1 year and 2 month wild-type (black) and Cx3Cr1-knockout (grey) microglia. **h)** Fold changes of regulatory matrisome genes that were significantly differentially expressed between 2-month Cx3Cr1 heterozygous (black) and Cx3Cr1-knockout (grey) microglia. **i)** Venn diagram depicting the proportion of matrisome-related genes that were uniquely impacted by age in wild-type and Cx3Cr1-knockout microglia (top).

**Extended Data Figure 5.**
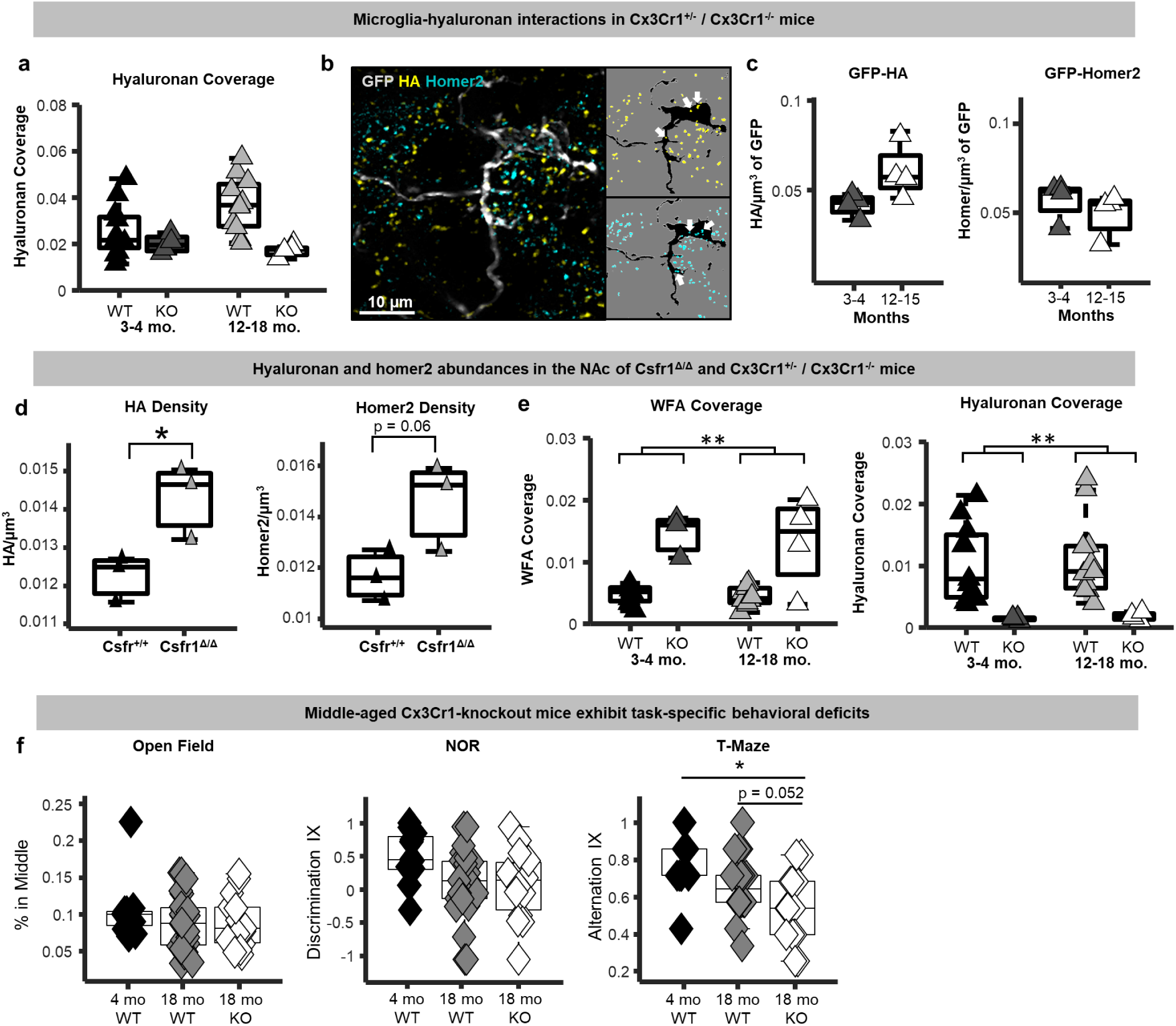
**a)** VTA hyaluronan tissue coverage in young-adult (3-4 months) and middle-aged (12-18 months) wild-type and Cx3Cr1-knockout (Cx3Cr1^EGFP/EGFP^) mice. **b)** Putative microglia engulfment of hyaluronan and homer2 was also examined in Cx3Cr1^EGFP/EGFP^ mice. **c)** Putative microglia-hyaluronan engulfment and microglia-homer2 engulfment in Cx3Cr1^EGFP/EGFP^ mice. **d)** NAc hyaluronan fibril and homer2 puncta densities in Csf1r^+/+^ (black triangles) and Csf1r^ΔFIRE/ΔFIRE^ (grey triangles) mice. There was a significant increase in hyaluronan abundance in the Csf1r^ΔFIRE/ΔFIRE^ mice compared to control (p < 0.05; unpaired t-test). **e)** WFA and hyaluronan tissue coverage in the NAc of young-adult (3-4 months) and middle-aged (12-18 months) wild-type and Cx3Cr1-knockout (Cx3Cr1^EGFP/EGFP^) mice. **f)** Performance of young-adult (4 months) wild-type, middle-aged wild-type (18 months), and middle-aged (18 months) Cx3Cr1-knockout mice on an open field test, a novel object recognition test, and a T-Maze test.

**Extended Data Figure 6.**
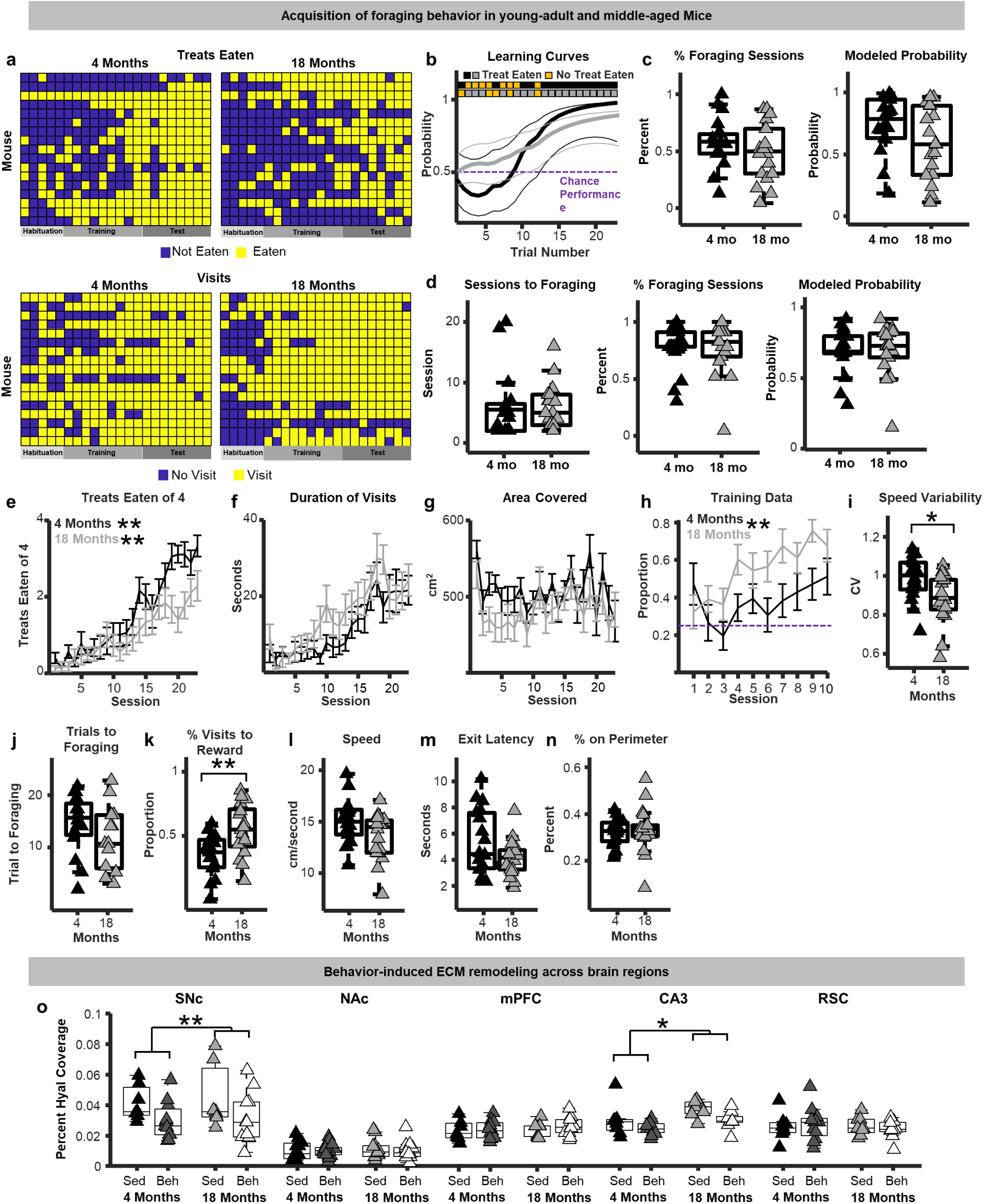
**a)** Behavioral raster plots of treats eaten (top) and feeder visits (bottom) during encoding across sessions used for hidden Markov modeling of foraging behavior. **b)** Representative learning curves from hidden Markov modeling for a young-adult and late-middle-aged mouse. **c)** The proportion of sessions mice exhibited foraging behavior (ate at least 1 treat), and modeled estimates of the probability of foraging using **c)** treat-eaten and **d)** feeder visits data for young-adult (4 months; black) and late-middle-aged (18 months; grey) mice. **e)** The number of treats eaten (of 4) across all encoding sessions for young-adult and middle-aged mice. Both young-adult and middle-aged mice exhibited increases in the number of treats eaten across foraging sessions (p < 0.01; n-way ANOVA). **f)** The duration of feeder visits also increased across sessions (p < 0.01; n-way ANOVA). **g)** The area covered across sessions for young-adult and late-middle-aged mice. **h)** The proportion of feeder visits to the correct feeder location during training sessions for young-adult and late-middle-aged mice. Middle-aged mice made more correct feeder visits during training compared to middle-aged (p < 0.01; n-way ANOVA). **i)** Variability in running speeds for young-adult and late-middle-aged mice. Middle-aged mice exhibited lower running speed variability compared to young (p < 0.05; unpaired t-test). **j)** The number of trials to consistent foraging behavior. **k)** The proportion of rewarded feeder visits across training sessions. A greater proportion of feeder visits were to the rewarded feeder in older mice (p < 0.01; unpaired t-test). **l)** Average running speed, **m)** the latency to exit the start box, and **n)** proportion of time on the perimeter of the arena. **o)** Boxplots depicting hyaluronan (hyal) tissue coverage in young-adult sedentary (black) and behavior-trained (dark grey) mice and late-middle-aged sedentary (light grey) and behavior-trained mice (white) in substantia nigra pars compacta (SNc), nucleus accumbens (NAc), medial prefrontal cortex (mPFC), CA3 region of the hippocampus, and retrosplenial cortex (RSC). Significant behavior-induced hyaluronan remodeling was observed in the SNc (p < 0.01; n-way ANOVA) and CA3 (p < 0.05; n-way ANOVA), but not in the NAc, mPFC, or RSC.

**Extended Data Figure 7.**
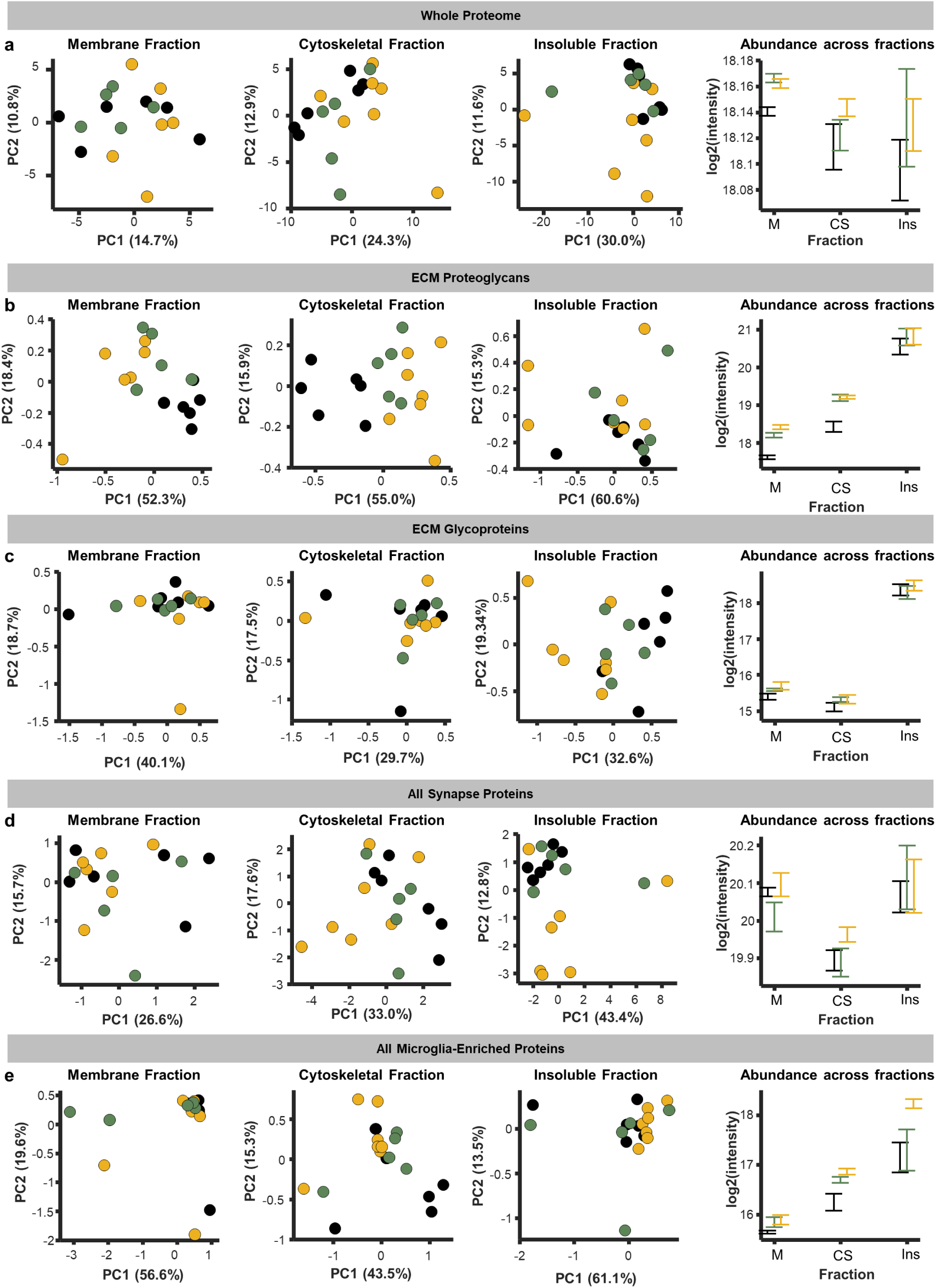
**a)** PCA plots of whole proteomes from membrane, cytoskeletal, and insoluble fractions (left 3 panels), and average log2-transformed protein intensities of the whole proteome across subcellular fractions. **b)** PCA plots of ECM proteoglycans from membrane, cytoskeletal, and insoluble fractions (left 3 panels), and average log2-transformed protein intensities of ECM proteoglycans across fractions. **c)** PCA plots of ECM glycoproteins from membrane, cytoskeletal, and insoluble fractions (left 3 panels), and average log2-transformed protein intensities of ECM glycoproteins across subcellular fractions. **d)** PCA plots of synapse proteins from membrane, cytoskeletal, and insoluble fractions (left 3 panels), and average log2-transformed protein intensities of synapse proteins across subcellular fractions. **e)** PCA plots of all microglia-enriched proteins from membrane, cytoskeletal, and insoluble fractions (left 3 panels), and average log2-transformed protein intensities of microglia-enriched proteins across subcellular fractions.

**Extended Data Figure 8.**
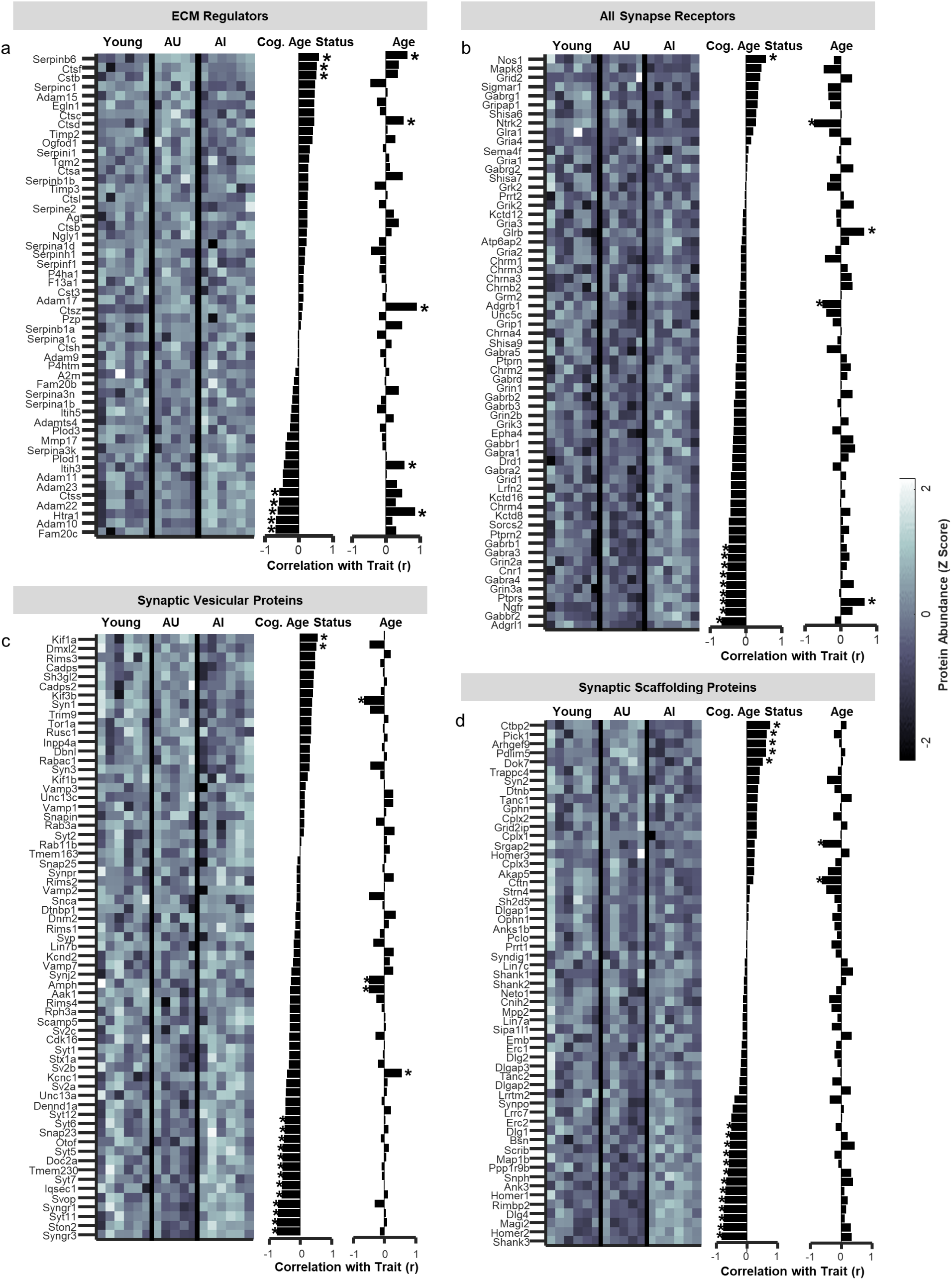
**a)** Heat plot of all detected ECM regulatory proteins abundances and bar plots of their relationship with cognitive status and age (* p < 0.05; linear probability model). **b)** Heat plot of all detected excitatory, inhibitory, and neuromodulator receptor abundances and bar plots of their relationship with cognitive status and age (* p < 0.05; linear probability model). **c)** Heat plot of all detected synaptic vesicle proteins abundances and bar plots of their relationship with cognitive status and age (* p < 0.05; linear probability model). **d)** Heat plot of all detected synaptic scaffolding proteins abundances and bar plots of their relationship with cognitive status and age (* p < 0.05; linear probability model).

**Extended Data Figure 9.**
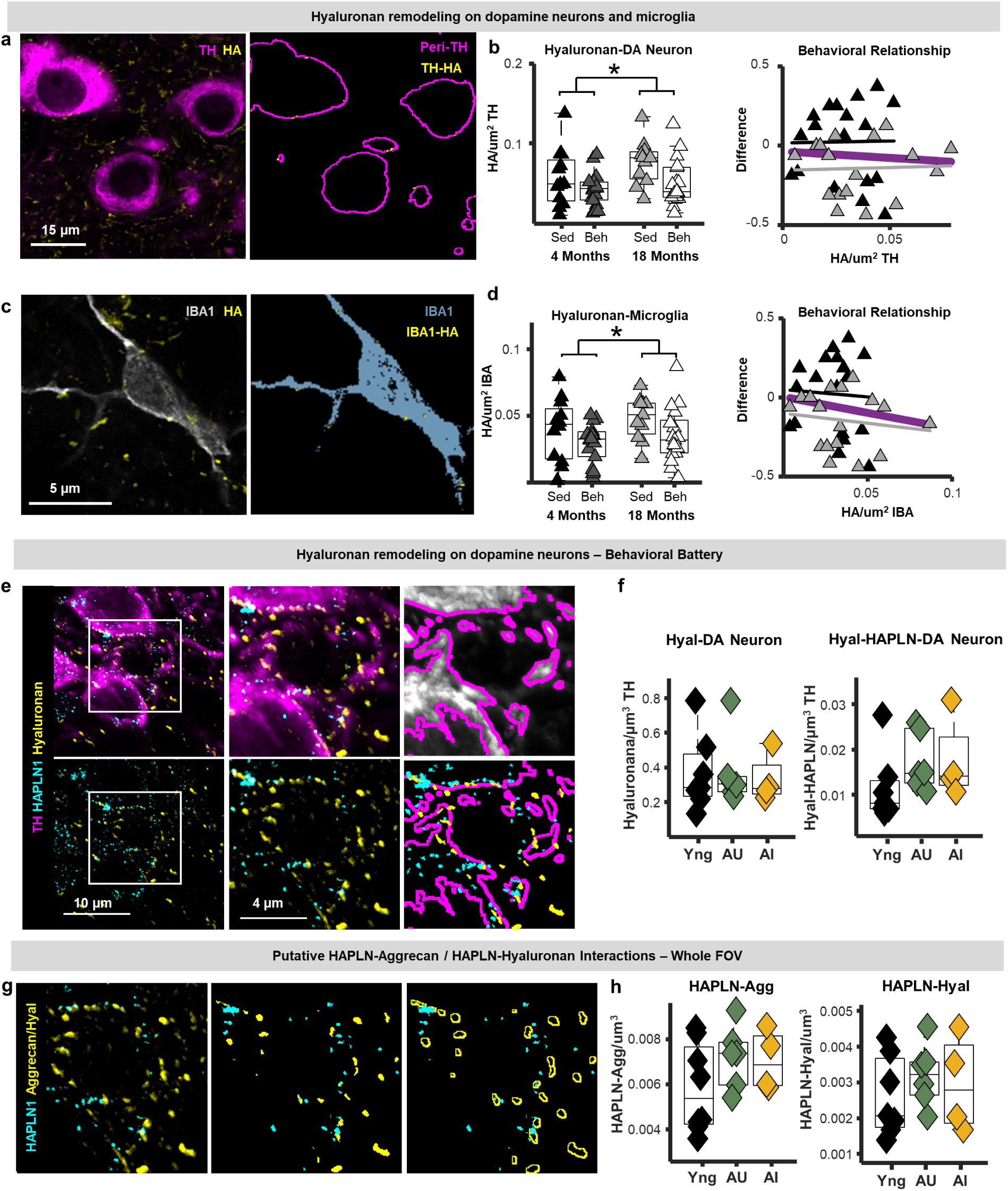
**a)** Left: example photomicrograph of histochemically labelled tyrosine hydroxylase (TH)-positive neurons and hyaluronan in the VTA of a middle-aged mouse. Right: peri-TH ROIs (magenta) used to index hyaluronan fragment on dopamine neurons (hyaluronan-DA neuron; yellow). **b)** Left: Boxplots depicting hyaluronan-DA neuron fibril densities normalized to the TH signal in young-adult sedentary (black) and behavior-trained (dark grey) mice and late-middle-aged sedentary (light grey) and behavior-trained mice (white). Behavior-trained mice exhibited lower hyaluronan-DA neuron densities relative to controls (p < 0.05; n-way ANOVA). Right: Relationship between TH-hyaluronan fibril densities and performance on reward-based foraging task (proportion of correct feeder visits difference measure). **c)** Left: example photomicrograph of an IBA1-positive microglia cell body and proximal branches and hyaluronan in the VTA of a middle-aged mouse. Right: IBA1 ROI used to examine hyaluronan fragments in contact with microglia (hyaluronan-microglia). **d)** Left: Boxplots depicting hyaluronan-microglia fibril densities normalized to the IBA1 signal in young-adult sedentary (black) and behavior-trained (dark grey) mice and late-middle-aged sedentary (light grey) and behavior-trained mice (white). Behavior-trained mice exhibited lower IBA1-hyaluronan densities relative to controls (p < 0.05; n-way ANOVA). Right: Relationship between IBA1-hyaluronan fibril densities and performance on reward-based foraging task (proportion of correct feeder visits difference measure). **e)** Example photomicrographs of histochemically labelled hyaluronan, hyaluronan and proteoglycan link protein 1 (HAPLN1), and tyrosine hydroxylase (TH) from the VTA of a late-middle aged mouse. The middle panels depict the fields of view depicted by the white squares in the left panel. Right: example of the peri-dopamine neuron region of interest used to estimate the abundance of hyaluronan and HAPLN1 on dopamine neuron surfaces in the VTA. **f)** Boxplots depicting hyaluronan-DA neuron puncta densities (left) and hyaluronan-HAPLN1-DA neuron puncta densities (right) in the VTA of young (black), aging unimpaired (AU; green), and aging impaired (AI; purple) mice. **g)** Schematic of hyaluronan-HAPLN and aggrecan-HAPLN1 proximity analysis to estimate the density of hyaluronan fibrils <0.5 microns from HAPLN1 without respect to the dopamine neuron surfaces. Note: this example photomicrograph is the same as in **e**. **h)** Boxplots depicting aggrecan-HAPLN1 (left) and hyaluronan-HAPLN1 (right) puncta densities (left) in the VTA of young, aging unimpaired (AU), and aging impaired (AI) mice.

## Acknowledgements

The authors would like to thank Keionna Newton and Fanny Etienne for providing tissue used in this study. Additionally, we thank Mia Donato and Claribel Charway for important contributions to mouse behavior pipelines used in this study. This work was supported by the American Federation on Aging Research grant number: MCKNIGHT21003, as well as by the Simons foundation Collaboration on Plasticity in the Aging Brain grant number: SFI-AN-NC-AB-Independence Postdoctoral-00007445.

## Author Contributions

**DTG** conceived the study, designed methodology, performed experiments, analyzed data, and wrote the manuscript. **AG** performed behavioral experiments and microglial analysis. **YJA** performed proteomic experiments and proteomic data analysis. **VP** performed proteomic experiments and proteomic data analysis. **LP** performed proteomic experiments comparing ECM enrichment strategies. **YZ** performed proteomic experiments comparing ECM enrichment strategies and provided critical support in manuscript preparation. **JAW** conceptualized proteomic experiments and analysis. **RAM** help conceptualize behavioral experiments and provided critical support in manuscript preparation. **LMD** conceived the study, designed methodology, and wrote the manuscript.

## Competing Interests

The authors do not claim any conflicts of interest. This research was supported in part by the Intramural Research Program of the National Institutes of Health (NIH). The contributions of the NIH author(s) were made as part of their official duties as NIH federal employees, are in compliance with agency policy requirements, and are considered Works of the United States Government. However, the findings and conclusions presented in this paper are those of the author(s) and do not necessarily reflect the views of the NIH or the U.S. Department of Health and Human Services.

## Notes

### Competing Interest Statement

The authors have declared no competing interest.

### Summary of Updates

This manuscript has incorporated novel histological and proteomic datasets in cognitively classified aging mice.

